# The differential metabolic signature of breast cancer cellular response to olaparib treatment

**DOI:** 10.1101/2022.06.14.495928

**Authors:** Domenica Berardi, Yasmin Hunter, Lisa van den Driest, Gillian Farrell, Nicholas J W Rattray, Zahra Rattray

## Abstract

Metabolic reprogramming and genomic instability are key hallmarks of cancer, the combined analysis of which has gained recent popularity. Given the emerging evidence indicating the role of oncometabolites in DNA damage repair and its routine use in breast cancer treatment, it is timely to fingerprint the impact of olaparib treatment in cellular metabolism. Here, we report the biomolecular response of breast cancer cell lines with DNA damage repair defects to olaparib exposure.

Following evaluation of olaparib sensitivity in breast cancer cell lines, we immunoprobed DNA double strand break foci and evaluated changes in cellular metabolism at various olaparib treatment doses using untargeted mass spectrometry-based metabolomics analysis. Following identification of altered features, we performed pathway enrichment analysis to measure key metabolic changes occurring in response to olaparib treatment.

We show a cell-line dependent response to olaparib exposure, and an increased susceptibility to DNA damage foci accumulation in triple-negative breast cancer cell lines. Metabolic changes in response to olaparib treatment were cell-line and dose-dependent, where we predominantly observed metabolic reprogramming of glutamine-derived amino acids and lipids metabolism.

Our work demonstrates the effectiveness of combining molecular biology and metabolomics studies for the comprehensive characterisation of cell lines with different genetic profiles. Follow-on studies are needed to map the baseline metabolism of breast cancer cells and their unique response to drug treatment. Fused with genomic and transcriptomics data, such readout can be used to identify key oncometabolites and inform the rationale for the design of novel drugs or chemotherapy combinations.

## INTRODUCTION

In a bid to develop new therapies against various cancer types, genomic instability, its underpinning mechanisms and contribution to tumorigenesis have been extensively investigated over the past few decades. Genomic instability, a well-known contributor to cancer, presents a therapeutic vulnerability that can be targeted in the development of novel chemotherapy agents (1).

To maintain their genomic integrity, cells are equipped with a range of DNA damage repair (DDR) pathways and responses to counteract DNA lesions formed in response to endogenous and exogenous insults (2). Hereditary mutations in these pathways have been correlated with increased cancer susceptibility, such that defects in homologous recombination contribute to approximately 10% of all breast cancers. These defects in DDR machinery result in the loss of function for genes implicated in DNA repair (i.e. breast cancer susceptibility gene 1/2- BRCA1/BRCA2) or dysregulation of cell cycle phases (3–5). While these genetic alterations increase the susceptibility to oncogenesis- they serve as therapeutic vulnerabilities-such that in the presence of a defective DNA repair pathway the inhibition of an alternate DDR mechanism will lead to cell death. This concept is referred to as synthetic lethality, which has formed the rationale for existing DDR inhibitors (6, 7). One such class of drugs, poly(ADP-ribose) polymerase (PARP) inhibitors, targets vulnerabilities in the homologous recombination DDR pathway (8).

PARP inhibitors as a class of DDR inhibitors block the activity of PARP enzymes involved in DNA damage repair; therefore, leading to accumulation of DNA double-strand breaks that gives rise to genomic instability and subsequent apoptosis (9). Several PARP inhibitors are currently approved as monotherapies for the treatment of locally advanced or metastatic breast cancer for patients with breast cancer harboring germline BRCA1/2 mutations or HER2- negative receptor status (8). In 2022, olaparib was approved by the FDA as an adjuvant treatment for patients with human epidermal growth factor receptor 2 (HER2)-negative and germline BRCA-mutated breast cancers following readout from the OlympiA trial (10).

While PARP inhibitors present a therapeutic opportunity for targeting DDR defects in breast and ovarian cancers, emerging evidence has shown a role for oncometabolites-small molecule intermediates of cellular metabolism- in determining the response to these chemotherapies. The biology of oncometabolites and their role in modulating DDR has been increasingly studied over the past few years, guiding new combination therapies and novel biological targets for drug discovery (1).

Metabolic reprogramming- a key feature of all cancers (11)- gives rise to chemoresistance in both treatment-naïve and treatment-resistant breast cancers (12). As with genomic instability, drivers of metabolic reprogramming can be broadly classified as intrinsic and extrinsic in origin (13). Intrinsic stimuli such as oncogenes and tumour suppressor genes, modulate cellular metabolism in breast cancer with several regulators including BRCA1/2, MYC, phosphatidylinositol-4,5-bisphosphate 3-kinase (PI3K) and p53 as examples. The functional interplay between these regulators of cellular metabolism, mediates DNA damage repair pathways and subsequent response to DDR chemotherapies. Recent evidence has shown that the upregulation of glucose utilization and glutamine metabolism are required to sustain increased tumour bioenergetic and biosynthetic demand, which vary according to the cellular genetic makeup (14). Intermediates from glucose and glutamine metabolism have been identified as key oncometabolites regulating the response to chemotherapy drugs, presenting novel biomarkers and potential actionable targets for novel drug discovery (13).

DDR mechanisms induce cellular metabolic changes through interference with purine and pyrimidine biosynthetic pathways, amino acid metabolism, protein biosynthesis and energy metabolism, impacting several metabolic routes (15). Mediators of DDR pathways, including PARP regulate several pathways exemplified by the pentose-phosphatase pathway, the TCA cycle and glycolysis. In breast cancer, PARP inhibition reduces glucose consumption and alters amino acid and nucleotide metabolism depending on the different cellular subtypes (16). Moreover, BRCA-1 deficient breast tumors appear to rely on glucose consumption through enhanced glycolysis (17). Differences in the metabolic signature between cell lines harboring different DNA repair mutations and measuring their response to PARP inhibitors can inform the rationale for selecting PARP inhibitors in certain breast cancer types and explore potential additional vulnerabilities as druggable targets (18).

DNA repair and regulation of metabolism are critical for maintaining homeostasis in normal human cells. However, the extensive dysregulation and aberrant function of both these pathways promotes tumorigenesis. Until recent, DNA repair and metabolic pathways have routinely been researched as distinct fields within their own right, but growing emerging research evidence an intrinsic inter-dependency between these pathways. Here, we report the differential cellular response of breast cancer cell lines with DDR defects to olaparib exposure through combined analysis of DNA damage and metabolomics profiling. Combined evaluation of the DNA damage response and metabolic reprogramming offers new opportunities in the development of novel chemotherapies against cancer.

## MATERIALS

### Cell lines and chemicals

All cell lines used in this study were purchased from the vendor and maintained in accordance with manufacturer instructions. All cell culture reagents were obtained from Gibco (Thermo Fisher Scientific). MCF7 (RRID:CVCL_0031, Sigma, EACC collection) and MDA-MB-231 cells (RRID:CVCL_0062, ATCC) were purchased and maintained in Dulbecco’s Modified Eagle Medium (DMEM, high glucose) supplemented with 10% *v/v* FBS (high glucose, Invitrogen), 1% *v/v* non-essential amino acids (NEAA) and 1% *v/v* penicillin-streptomycin (Invitrogen). Corresponding cell line origins, hormone receptor status and mutational profiles are included in **Table S 1**. HCC1937 cells obtained from ATCC (RRID:CVCL_0290) were maintained in RPMI supplemented with 10% *v/v* FBS and 1% v/v penicillin-streptomycin. All cell lines were maintained at 37 °C in a pre-humidified atmosphere containing 5% *v/v* CO_2_ and used within ten passages for the purposes of this work (passage 2-10). Olaparib (SantaCruz Biotechnology Inc.) was prepared as a 100 mM stock solution in DMSO, aliquoted and stored at -20 °C until use. γH2AX, p53BP1 primary antibodies (Cell Signalling Technologies) were used for foci immunostaining alongside the Alexa Fluor^®^ 488-conjugated secondary antibody (Fisher Scientific).

## METHODS

### Cell Viability Assays

MCF-7, MDA-MB-231 and HCC1937 cells undergoing exponential growth were seeded at a density of 4,000 cells/well in 96 well plates and incubated overnight to facilitate cell attachment. On the following day, cells were exposed to either blank growth medium (control) or growth medium containing different concentrations of olaparib (treatment medium) ranging from 0.01-500 µM for seven days at 37 °C and 5% *v/v* CO_2_. Treatment media were replaced every three days with treatment medium. Following a seven-day incubation, cell viability was measured using CellTiter 96^®^ Aqueous Non-Radioactive Cell Proliferation Assay (Promega) (3-(4,5-dimethylthiazol-2-yl)-5-(3-carboxymethoxyphenyl)-2-(4-sulfophenyl)-2H-tetrazolium (MTS) reagent. The resultant absorbance at 490 nm was measured using a GM3500 Glomax® Explorer Multimode Microplate Reader (Promega).

Growth curves represent percentage cell growth following treatment with different concentrations of olaparib and are plotted as a semi-log dose-response curve. The half maximal inhibitory concentration (IC_50_) was determined using a linear regression model. Statistical analysis was performed using GraphPad Prism (RRID:SCR_002798, v.9.0.1). Three independent biological replicates (five wells per treatment concentration) were performed for each cell line.

### Immunostaining for γH2AX and p53BP1

Foci immunodetection for γH2AX and p53BP1 was performed in both control (growth medium) and for cells treated with olaparib (IC_10_, IC_25_ and IC_50_ doses) for seven days. Briefly, cell monolayers were fixed in chilled 4% *w/v* formaldehyde containing 2% *w/v* sucrose in PBS, followed by fixation in ice-cold methanol (100% v/v). Subsequently, cells were permeabilized in 0.25% *v/v* Triton X-100 in PBS, blocked with 5% *v/v* goat serum/5% *w/v* BSA, immunoprobed with either a primary rabbit anti-γH2AX antibody (RRID:AB_420030) (1:1000) or primary rabbit anti-P53BP1 (1:200) antibody (RRID:AB_11211252, CST #2675 for p53BP1) overnight at 4 °C. Cell monolayers were treated with goat, anti-rabbit Alexa Fluor^®^ 488 conjugated secondary antibody and counterstained with DAPI. Image acquisition was carried out using an Invitrogen EVOS Auto Imaging System (AMAFD1000-Thermo Fisher Scientific) with a minimum of 100 cells imaged per treatment condition. Resultant foci images were analysed in Cell Profiler (v.4.2.1.) using a modified version of the speckle counting pipeline.

### Sample preparation and metabolite extraction

MCF-7, MDA-MB-231 and HCC1937 cells were seeded at a density of 2 x 10^6^ cells per well in 6-well plates, and exposed to growth medium containing olaparib at IC_10_, IC_25_ and IC_50_ doses, as determined from the MTS assay (n=5 per treatment concentration). Following exposure to olaparib, the growth medium was aspirated from each well, centrifuged to remove cell debris, and stored at -80 °C. Next, treated cells were washed with pre-chilled PBS, with the metabolites quenched and extracted in a final volume of 1.5 mL pre-chilled (-80°C) mixed solvent (Methanol:Acetonitrile:Water=50:30:20). Resultant cell pellets were collected, and submerged in liquid nitrogen, vortexed and sonicated for 3 minutes in an ice-water bath. This procedure was performed in triplicate. Resultant extracts were centrifuged at 13,000x g for 10 minutes at 4 °C and the pellets were retained for protein quantification using the Bradford assay. The resultant supernatant was collected, and dried with a Speed vac centrifuge (Savant-SPD121P). Dried metabolite pellets were reconstituted in Acetonitrile:Water (50:50) at volumes normalized to the relative protein content. Quality control (QC) samples were prepared by pooling samples across all control and treatment groups. Solvent blank and QC samples were inserted in analytical batch after every five samples to assess the stability of detecting system.

### Liquid Chromatography Tandem Mass Spectrometry (LC-MS/MS)

Metabolite separation was performed on a binary Thermo Vanquish ultra high performance liquid chromatography system where 5µl of reconstituted cellular extract was injected on to a Thermo Accucore HILIC column (100mm x 2.1 mm, particle size 2.6 µm). The temperature of the column oven was maintained at 35 °C while the autosampler temperature was set at 5 °C. For chromatographic separation, a consistent flow rate of 500 µL/min was used where the mobile phase in positive heated electrospray ionisation mode (HESI+) was composed of buffer A (10 mM ammonium formate in 95% acetonitrile, 5% Water with 0.1% formic acid) and buffer B (10 mM ammonium formate in 50% acetonitrile, 50% Water in 0.1% formic acid). Likewise, in negative ionisation mode (HESI-) buffer A (10 mM ammonium acetate in 95% acetonitrile, 5% water with 0.1% acetic acid) and buffer B (10 mM ammonium acetate in 50% acetonitrile, 50% water with 0.1% acetic acid). The elution gradient used for the chromatographic separation of metabolites is included in supplementary information (**Table S2**).

A high-resolution Exploris 240-Orbitrap mass spectrometer (ThermoFisher Scientific) was used to perform full scan and fragmentation analyses. Global operating parameters were set as follows: spray voltages of 3900 V in HESI+ mode, and 2700 V in HESI-mode. The temperature of the transfer tube was set as 320 °C with a vaporiser temperature of 300 °C. Sheath, aux gas and sheath gas flow rates were set at 40, 10 and 1 Arb, respectively. Data dependent acquisitions (DDA) were performed using the following parameters: full scan range was 70 – 1050 m/z with a MS1 resolution of 60,000. Subsequent MS/MS scans were processed with a resolution of 15,000. High-purity nitrogen was used as nebulising and as the collision gas for higher energy collisional dissociation. Further details are included in supplementary information.

### Mass Spectrometry Data Processing

Raw data files obtained from Thermo Scientific Xcalibur^TM^ software 4.2 were imported into Compound Discoverer^TM^ 3.2 software where the “Untargeted Metabolomics with Statistics Detect Unknowns with ID Using Online Databases and mzLogic” feature was selected (supplementary information). The workflow analysis performs retention time alignment, unknown compound detection, predicts elemental compositions for all compounds, and hides chemical background (using Blank samples). For the detection of compounds, mass and retention time (RT) tolerance were set to 3 ppm and 0.3 min, respectively. The library search was conducted against the mzCloud, Human Metabolome Database (HMDB) and Chemical Entities of Biological Interest (ChEBI) database. A compound table was generated with a list of putative metabolites (known and unknown). Among them, we selected all the known compounds fully matching at least two of the annotation sources. The selected metabolites were then used to perform pathway and statistical analysis.

### Pathway Analysis with MetaboAnalyst

Prior to analysis of the metabolic pathways with MetaboAnalyst 5.0 (RRID:SCR_015539, https://www.metaboanalyst.ca/), a HMDB identification code was assigned to each selected metabolite. A joint pathway analysis was performed by integrating the genes relative to each cell line (Table 1) with the list of ID compounds and their associated Log2 Fold change values. The integration method combined both genes and metabolites into a single query, then used to perform the enrichment analysis. This latter was based on a hypergeometric test. Finally, important nodes (compounds) were scored based on their betweenness centrality, and pathway analysis results were generated.

### Statistical Analysis

All data are presented as mean ± standard deviation (n≥5). For metabolomics analysis, Principal Component Analysis (PCA) was performed to test analytical reproducibility of QC injections, reduce the dimensionality of our data and determine the metabolic profiles of the different sample groups. Differential analysis was used to compare differences between control and treatment groups and plotted as a Volcano plot (log-fold change vs. -log10 p-value). Peak areas were log_10_ transformed and p values calculated for the sample group by analysis of variance (ANOVA) test. A p value<0.05 and fold-change of 1.5 was deemed to be statistically significant.

## RESULTS

### Olaparib sensitivity analysis

To determine the olaparib dose range for subsequent foci and metabolomics experiments, we measured the sensitivity of MCF7, MDA-MB-231 and HCC1937 cell lines to olaparib exposure over a seven day treatment duration. The rationale behind exploring sensitivity to olaparib in these cell lines, was to perform a comparison between two triple-negative (MDA-MB-231 and HCC1937) and a non-triple-negative (MCF-7) cell line.

Our results show that exposure to olaparib caused a reduction in cell viability in all cell lines in a dose-dependent manner (**Figure 1**). We observed superior efficacy of olaparib in reducing cell viability in both MCF7 and MDA-MB-231 cells, with a calculated half maximal inhibitory concentration (IC_50_) of 10 µM and 14 µM, respectively. However, in the case of HCC1937 cells, a higher concentration of olaparib was required to achieve the same reduction in cell viability (150 µM), indicating a lower efficacy of response to olaparib in this cell line.

**Figure 1.**
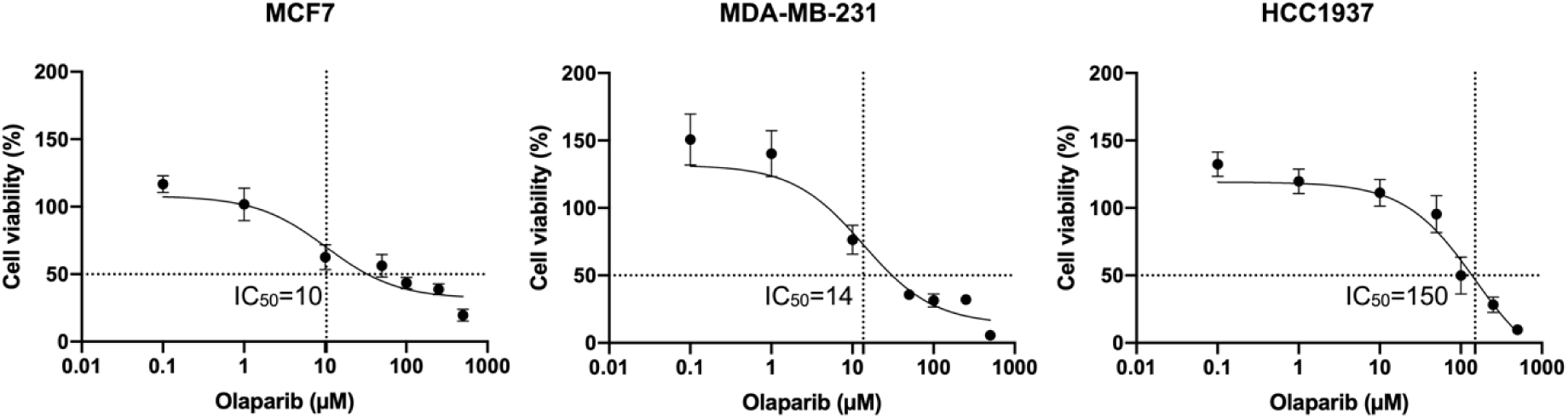
Corresponding MTS dose-response curves for MCF7, HCC1937 and MDA-MB-231 cells treated with ascending doses of olaparib (0.1-500 µM) for seven days. The corresponding R^2^ values for fitted dose-response curves in MCF7 (IC_50_= 10 µM), MDA-MB-231 (IC_50_= 14 µM), and HCC1937 (IC_50_= 150 µM) cells were 0.89, 0.91 and 0.85, respectively.

### Exposure to olaparib induces dose-dependent formation of γH2AX and 53BP1 foci in breast cancer cells

PARP inhibition induced by olaparib exposure results in the accumulation of DNA damage in cells by compromising their DDR mechanisms. Therefore, we next investigated the extent to which olaparib exposure at various doses (IC_10_, IC_25_ and IC_50_- determined from MTS assays) promotes the accumulation of DNA double strand breaks (DSBs) in MCF-7, MDA-MB-231 and HCC1937 cell lines. Key markers for DNA DSB formation include phosphorylated histone H2 variant H2AX (γH2AX) (19) and the damage sensor p53-binding protein 1 (p53BP1), which are rapidly recruited to sites of DNA damage and their accumulation is directly proportional to the number of DSB lesions (20). To measure the extent of DNA DSB formation, we performed immunofluorescence of p53BP1 and γH2AX foci.

Based on our results, p53BP1 and γH2AX foci levels increased in a dose-dependent manner in both MCF7 and MDA-MB-231 cells in response to ascending doses of olaparib (**Figure 2a, b,d,e**; **Figure 3a,b,d,e**). However, in HCC1937 cells, the expression of both foci decreased at the highest olaparib treatment dose (150 µM), in comparison to the 15 and 50 µM exposure doses (**Figure 2c, f**; **Figure 3c,f**). Generally, a higher number of both p53BP1 (mean >10 foci per cell) and γH2AX (mean > 20 foci per cell) foci were observed in the HCC1937 cell line, compared to the MCF7 and MDA-MB-231 cells, where a mean of <10 foci per cell were measured for both markers. These results are consistent with the dose-dependent sensitivity of MCF7 and MDA-MB-231 cells in response to olaparib exposure, further confirming cell-line dependent response to olaparib exposure.

**Figure 2.**
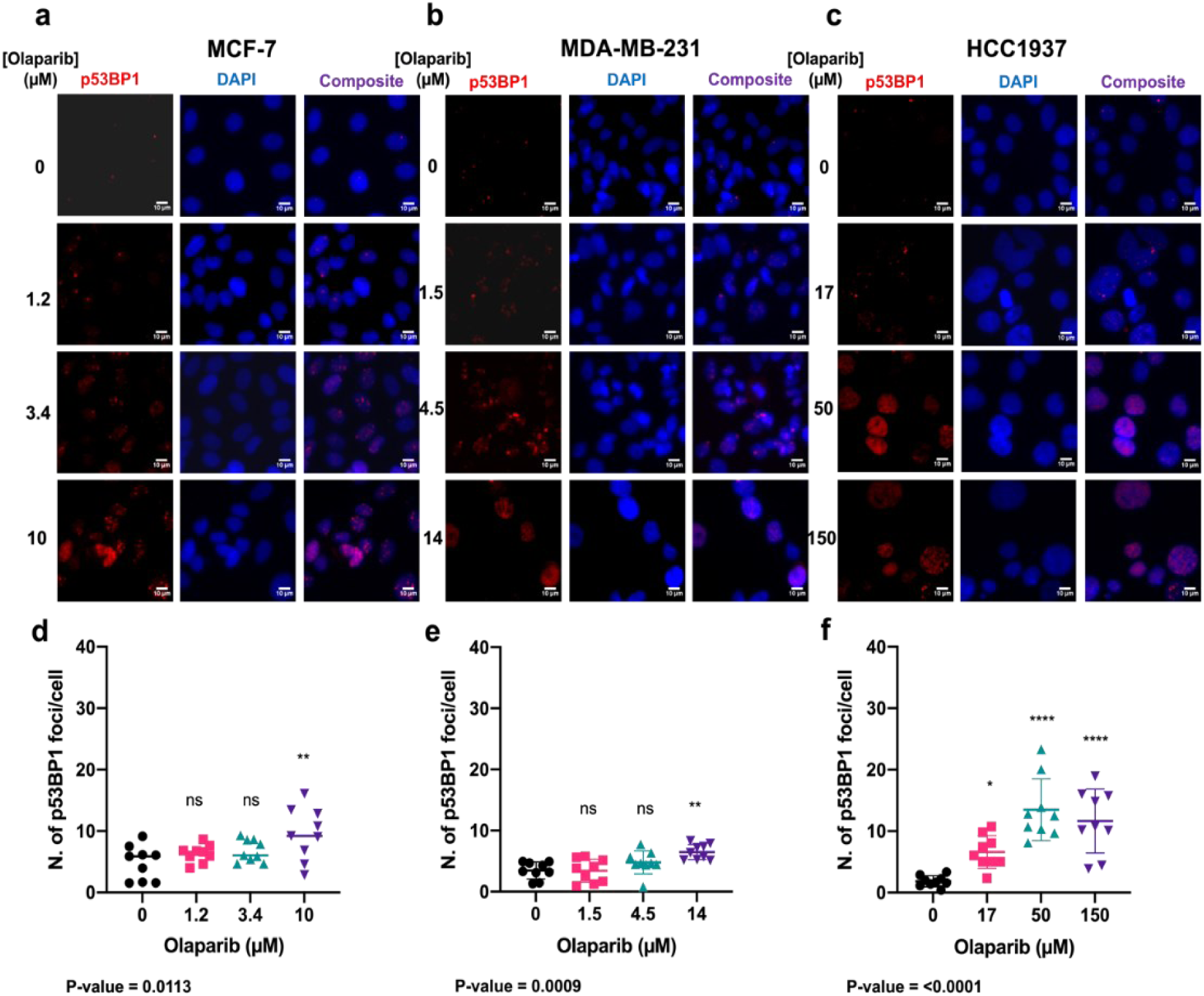
The formation of p53BP1 foci in response to treatment with either growth medium or medium containing olaparib. Representative images of immunolabelled P53BP1 foci (red), DAPI (blue) nuclear counterstain and composite (p53BP1 (red) and DAPI (blue)) in MCF-7, MDA-MB-231, and HCC1937 cells treated with olaparib for seven days (a-c). Corresponding p53BP1 foci counts determined using Cell Profiler (d-f). 9 repeats with on average >100 cells per each sample. p-values have been determined through ANOVA test. Dunnett’s multiple comparison test was used as a follow up to ANOVA test and the p-values were represented as: non-significant=ns, 0.05=*, 0.005=**, 0.0005=***, >0.00005=****.

**Figure 3.**
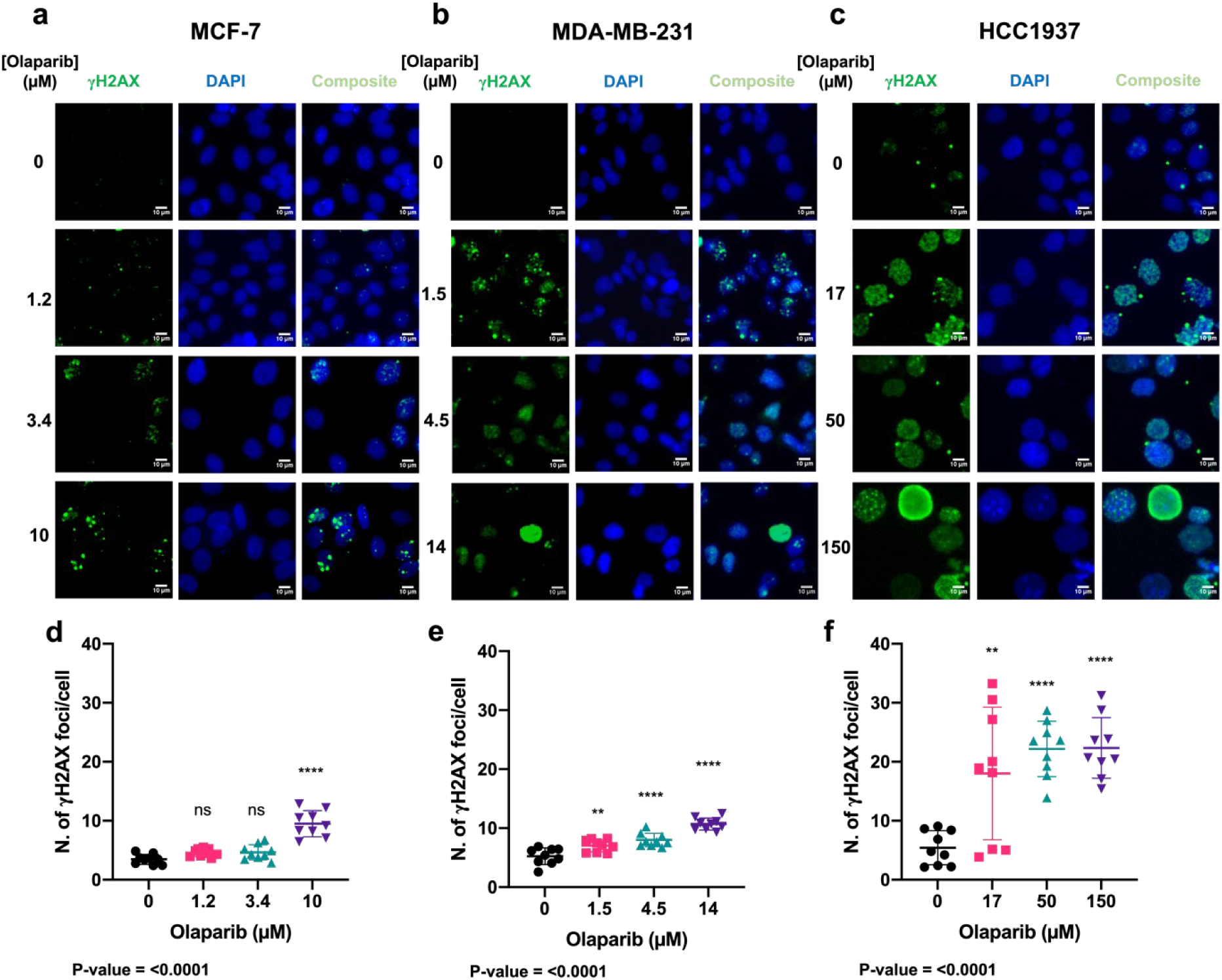
The formation of γH2AX foci formation in response to treatment with either growth medium or medium containing olaparib. Representative images of immunolabelled γH2AX foci (green), DAPI (blue) nuclear counterstain and composite (γH2AX and DAPI) in MCF-7, MDA-MB-231, and HCC1937 cells treated with for seven days (a-c). Corresponding γH2AX foci counts determined using Cell Profiler (d-f). (>100 cells per sample). Dunnett’s multiple comparison test was used as a follow up to ANOVA and corresponding p-values were represented as: non-significant=ns, 0.05=*, 0.005=**, 0.0005=***, >0.00005=****.

### Biomolecular pathways altered in response to olaparib exposure vary across different cell lines

To comprehensively measure the extent of variation induced by olaparib exposure in MCF-7, MDA-MB-231 and HCC1937 cell lines, we profiled their metabolome using an in-house untargeted liquid chromatography-mass spectrometry-based metabolomics pipeline (**Figure 4A**). After data acquisition, data processing and analysis were performed in Compound Discoverer 3.2. First, we used principal component analysis (PCA) to visualise and interpret the clustering of quantified metabolite data to examine global differences between treatment groups and cell lines examined, which was followed by pairwise PCA between control and treated groups across positive and negative analysis modes (**Figure 4**).

**Figure 4.**
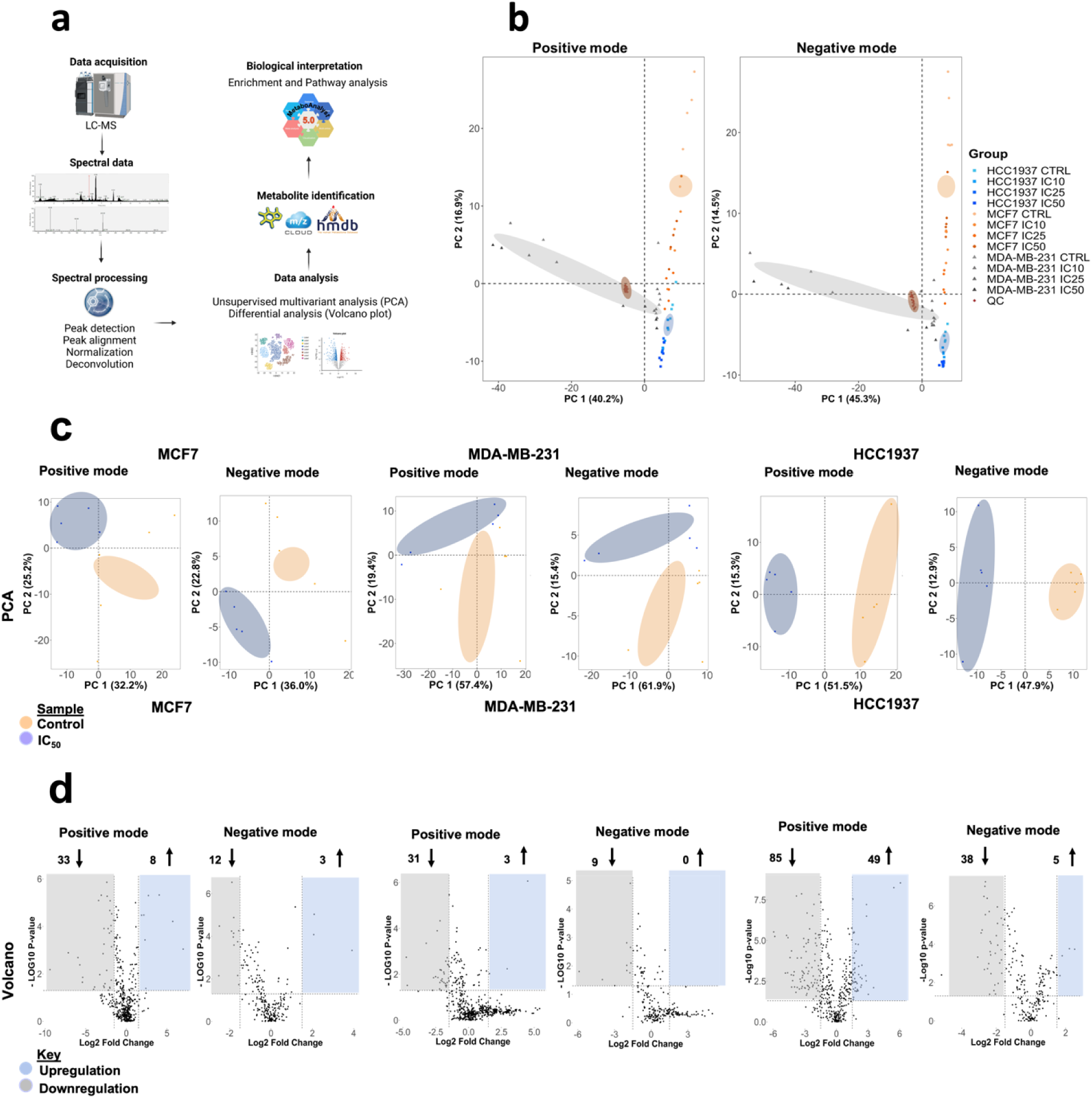
Statistical analyses of global metabolic features identified in MCF7, MDA-MB-231 and HCC1937 upon exposure to IC_10_, IC_25_ and IC_50_ olaparib doses for seven days acquired in positive and negative ionization mode. a) Workflow used in this study to perform pathway analysis from metabolomics analyses. b) Global PCA score plots of the analysed breast cancer cell lines for data acquired in positive and negative ionization mode. For each treatment group, five replicates were used. Data points in the two-dimensional PCA score plot were central scaled. c) PCA pairwise analysis and differential analysis of metabolites altered in IC_50_- treated cells, d) Volcano plots displaying enriched (blue) and depleted (grey) metabolic features by representing the log2 fold change in altered features and the -log10 adjusted p-values with cut off values selected at >1.5 and <0.05, respectively.

Pooled quality control (QC) data confirm the stability of the data acquisition system across all the measurements performed in positive and negative ionization acquisition modes (**Figure 4 b**). Distinct clustering patterns were observed, with better separation for the IC_50_ olaparib treatment dose across all cell lines (**Figure 4c**, **Figure S1**). Volcano plots indicate the differential number of metabolic features that are significantly altered following exposure to olaparib, relative to control (**Figure 4d, Figure S2**). From a metabolic perspective, we observed that HCC1937 (BRCA1-mutated) cells were the most susceptible to exposure at the IC_50_ olaparib treatment dose, while the MCF7 cells showed a higher number of significantly altered metabolic features at the IC_25_ olaparib treatment concentration (**Table S4**). Together, these findings show a differential dose- and cell line-dependent metabolic response to olaparib exposure.

### Amino acid and lipid metabolism are significantly altered in response to olaparib exposure

To analyse specific biomolecular pathways altered by olaparib exposure, we used MetaboAnalyst to identify key metabolic pathways significantly perturbed by olaparib treatment, and performed enrichment analysis for both control and treated samples (**Figure 5**, **Figure S3**). Among the pathways ranked in the top ten, we selected altered pathways with a corresponding pathway impact >0.1, and a p-value <0.05. (**Table S5**).

**Figure 5.**
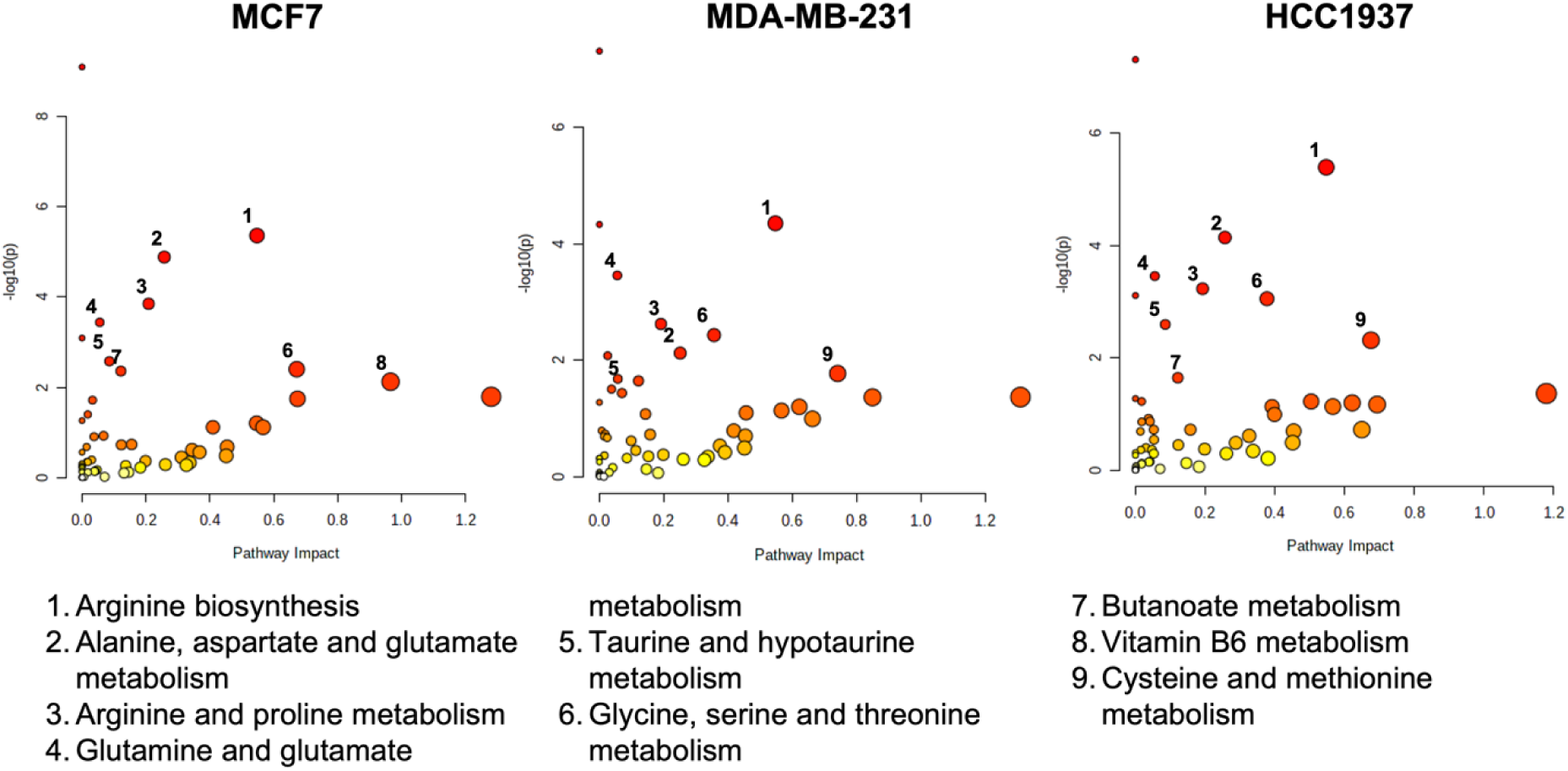
Pathway enrichment analysis of MCF7 (10 µM), MDA-MB-231 (14 µM) and HCC1937 (150 µM) cells following a seven-day exposure to olaparib. Enrichment analysis was based on the hypergeometric test. Topological analysis was based on betweenness centrality. The tight integration method was used by combining genes and metabolites into a single query. A p<0.05, and pathway impact >0.1 were deemed significant.

Across all cell lines examined, the top ten putative pathways significantly altered in Metaboanalyst (**see Figure 5**) were based on amino acid (arginine biosynthesis, glutamine, glycine, serine and threonine metabolism) and lipid metabolism (butanoate metabolism). Following the identification of metabolic pathways altered by olaparib exposure, we constructed a Venn diagram (**Figure S4**) to outline common overlapping and cell line-specific altered metabolic features.

Overlapping pathways are mostly represented by amino acid metabolism (glutamine, glutamate, aspartate, alanine, arginine and proline), suggesting a strong reliance of breast cancer cell metabolism on amino acids under baseline conditions (control samples). Upon olaparib exposure, the same pathways (amino acid metabolism) were among the most significantly-altered across all cell lines, while fatty acid (butanoate metabolism) and vitamin B6 metabolism were only significantly perturbed in MCF-7 cells.

Next, we explored individual metabolites that were associated with significantly altered metabolic pathways in response to olaparib exposure and evaluated relative changes in their levels between control and treatment samples. These results are presented through a heatmap clustering analysis (**Figure 6**). A correlation analysis between each metabolite is shown in **Figure S5**, and a wider list of compounds specific for each cell type is provided in **Table S6**.

**Figure 6.**
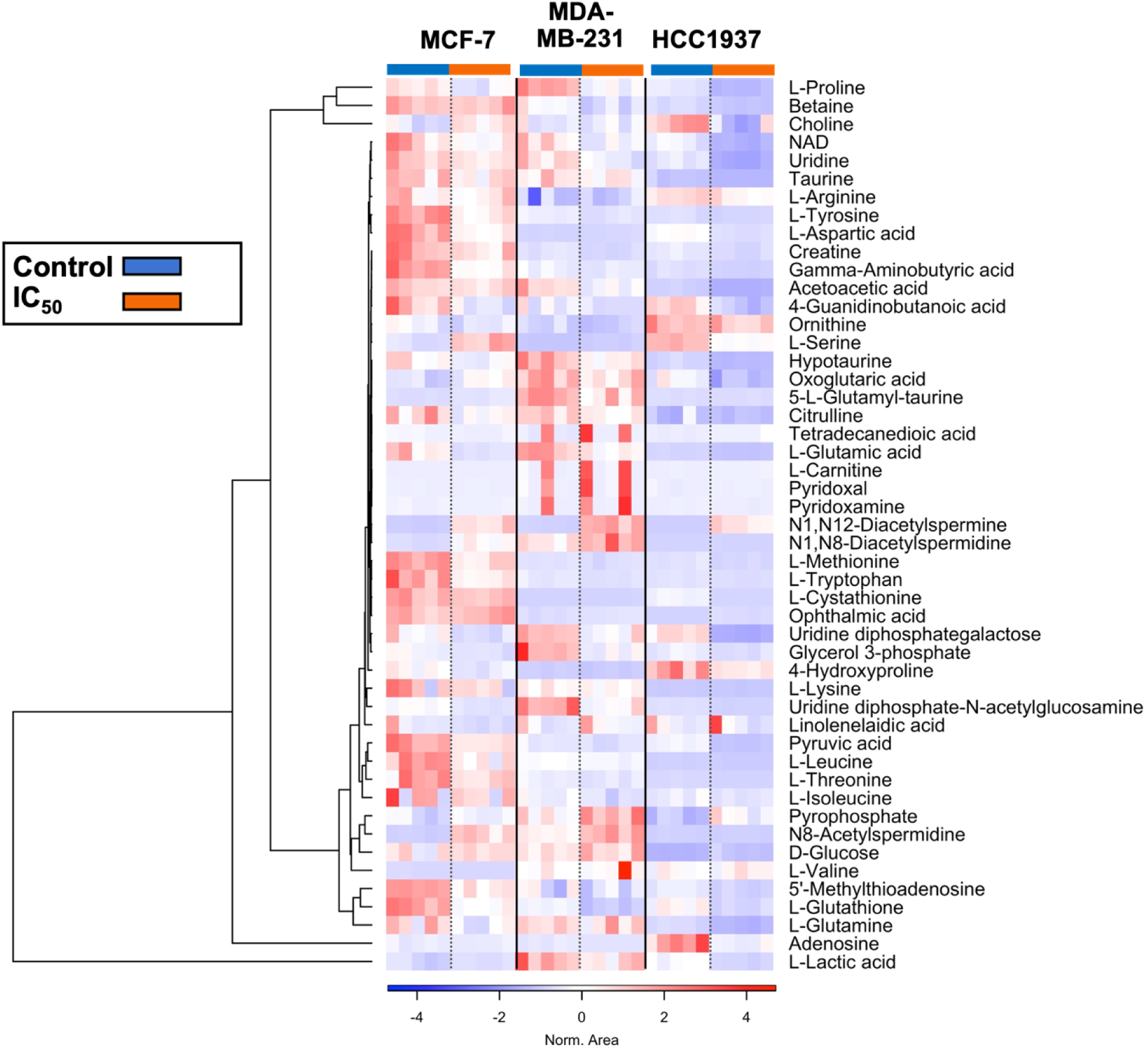
Heatmap cluster analysis of relevant metabolites associated with the pathways altered upon exposure to olaparib in MCF7 (10 µM), MDA-MB-231 (14 µM) and HCC1937 (150 µM) cells for seven days. Clustering and distance function are Ward and Euclidean, respectively. Normalised areas indicate chromatographic peaks areas that have been normalised based on the QC samples to compensate for batch effects.

Multiple amino acids (glutamine, glutamate, arginine, proline, methionine, glycine, threonine, taurine, and hypotaurine) were found to be depleted following olaparib exposure (relative to control) in all cell lines examined. Arginine and proline metabolism were significantly depleted by olaparib exposure, with depletion of their derived polyamines detected in all cell lines examined. Conversely, catabolic products of arginine and proline metabolism (N8- Acetylspermidine, N1-N8-Diacetylspermidine, and N1-N12-Diacetylspermine) were enriched. Elevated levels of serine were observed in MCF7 and MDA-MB-231 cells, while depletion of serine levels was seen in HCC1937 cells.

Alpha-ketoglutarate (α-KG- glutamine-derived intermediate of the TCA cycle) was enriched in MCF7 and depleted in MDA-MB-231 and HCC1937 cells. A negative correlation was observed between α-KG and glutamine levels, and a positive correlation between α-KG, and citric and fumaric acid (TCA cycle intermediates). Aspartate (a TCA cycle product), accumulated in the KRAS-mutant MDA-MB-231 cells, while aspartate depletion was observed in MCF7 and HCC1937 cells. Glucose levels were significantly elevated relative to control samples in HCC1937 cells. Asparagine (a byproduct of aspartate) was absent in MDA-MB-231 cells, while its enrichment was detected in MCF7 and HCC1937 cells. In parallel, accumulation of AMP was observed in both MCF7 and HCC1937 cell lines, while it was absent in MDA-MB-231 cells and enrichment of PPi was detected in all cell lines examined following olaparib exposure.

In the case of lipid metabolism, we observed a global depletion of phosphocholines (PC) and phosphoethanolamines (PE) in all cell lines following olaparib treatment. Acylcarnitine levels varied across the cell lines, with an overall enrichment of long (C14 – C21) and very-long chain acylcarnitines (>C22) in all cell lines treated with olaparib. Moreover, we observed enriched alpha-linoleic acid (a polyunsaturated fatty acid-PUFA) levels in MCF7 and MDA-MB-231 cells, which was absent in HCC1937 cells.

Compared to non-treated cells, elevated levels of glucose were detected in all cell lines studied following olaparib treatment, while downregulation of most nucleobases was observed. Finally, NAD+ downregulation was detected in all cell lines treated with olaparib.

An overview of the metabolic features altered in response to olaparib exposure is given in **Figure 7**, where we mapped cell line differences in metabolite levels through the Kyoto Encyclopaedia of Genes and Genomes (KEGG) database.

**Figure 7.**
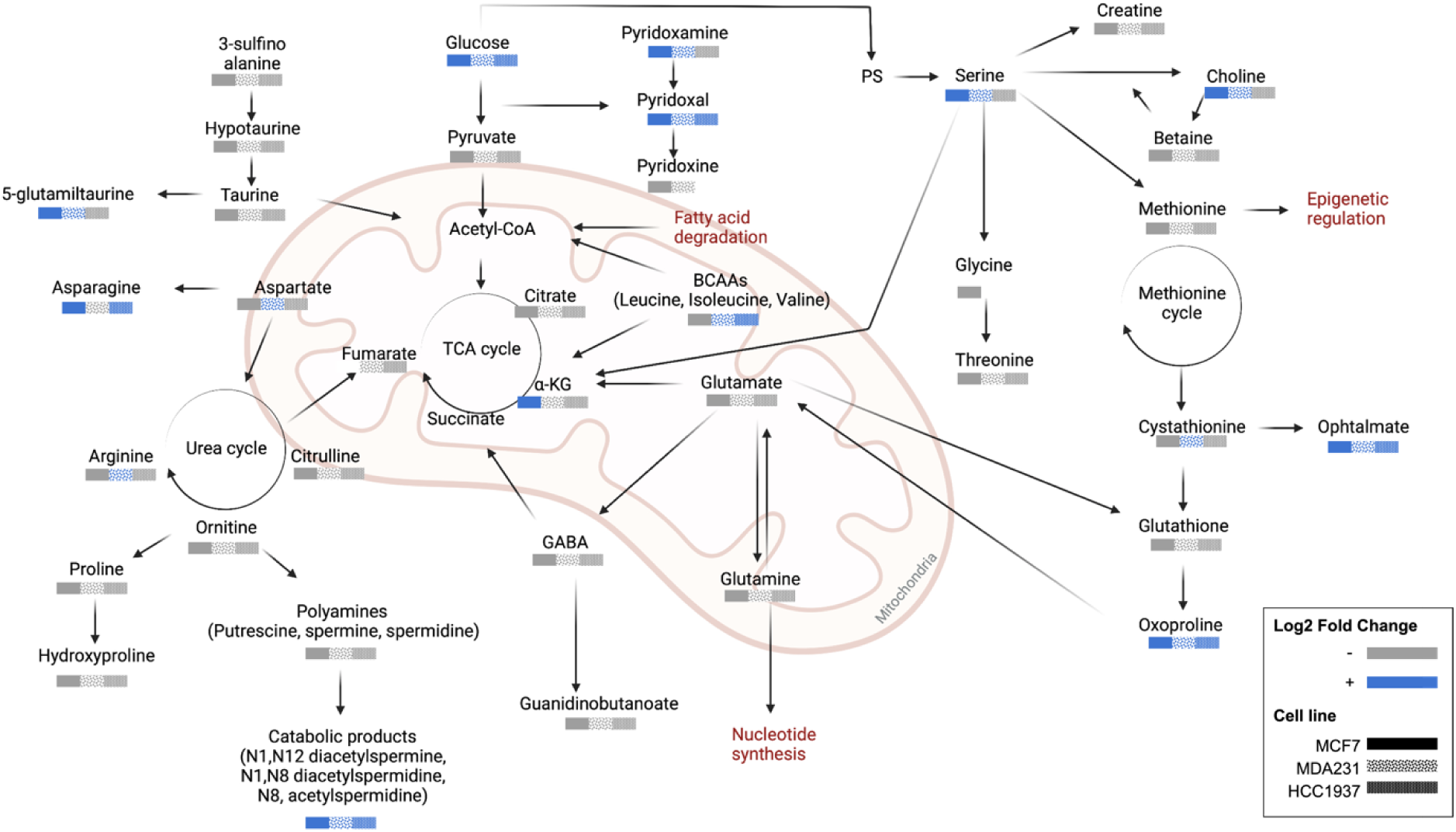
A summary of putatively identified metabolic pathways altered in response to olaparib exposure. Significantly altered features with a Log2 fold change of >1.5 (blue-enriched and grey-depleted). MCF-7 ( ), MDA-MB-231 ( ), and HCC1937 ( ).

## DISCUSSION

PARP inhibitors have shown promising results in the treatment of metastatic breast cancers harbouring germline BRCA1/2 mutations (21, 22). Recent clinical studies have shown evidence of PARP inhibitor efficacy in the management of breast cancer, irrespective of tumour BRCAness. Prior work has shown that BRCA1-mutated cells carrying a TP53 mutation are resistant to treatment with PARP inhibitors (23). Therefore, additional factors beyond BRCAness may govern sensitivity to PARP inhibition.

In this study we analyzed the sensitivity of two triple-negative (MDA-MB-231 and HCC1937) and MCF-7 (ER+, PR-, HER2-) cell lines to olaparib PARP inhibition (PARP1/2). The rationale for selecting these cell lines was to explore how their different genetic profiles define the observed differential biomolecular perturbations in response to olaparib treatment. Initially, we examined the responsiveness of MCF-7, MDA-MB-231 and HCC1937 cell lines to olaparib exposure using the MTS cell viability assay (**Figure 1**). Our results show differential sensitivity to olaparib exposure across the cell lines examined, with MCF-7 and MDA-MB-231 showing sensitivity to olaparib treatment at lower micromolar concentrations, and the BRCA1-mutant HCC1937 cell line showing less sensitivity (IC_50_- 150 µM). These findings are in agreement with previous reports of HCC1937 resistance to PARP inhibition, where the identification of predictive biomarkers of response to PARP inhibitor treatments was recommended beyond BRCA1/2 status (23).

Our analysis of γH2AX and p53BP1 DNA DSB immunolabelled foci (**Figure 3**) showed a higher occurrence of DNA damage foci in HCC1937 cells in comparison with MCF-7 and MDA-MB-231 cells with wild-type BRCA status. These observations suggest that BRCA status does not necessarily translate to olaparib sensitivity, and additional DDR components may define responsiveness. At present, routine clinical decision making surrounding the selection of treatment interventions are based on BRCA status, anatomical location, hormone receptor status and tumour stage, with very limited attention given to other mediators of DDR- namely homologous recombination-known to confer a BRCAness phenotype similar to BRCA 1 or 2 loss. Several recent studies have used whole-genome sequencing or the integration of homologous recombination panel scoring systems to provide an additional framework for predicting responders to PARP inhibitor treatment (24, 25).

Genetic biomarkers are routinely used in the clinical stratification of breast cancers and predicting treatment-emergent resistance (26). While genome-wide studies have improved patient stratification efforts, they lack the potential to account for functional phenotypic effects resulting from protein expression levels, or gain- or loss of function effects. Metabolomics has emerged in the past decade as an additional research toolbox for studying potential biomarkers of breast cancer with a range of applications ranging from early detection to the discovery of new metabolites and prognostic classification of patients with breast cancer (27).

Our goal in the present study was to apply combined analysis of DNA damage foci formation with global untargeted mass-spectrometry based metabolomics to map the metabolic changes occurring following exposure to olaparib. We examined the baseline differences in cellular metabolism across the cell line panel and extended this evaluation to examine cell line dependent response to olaparib treatment. Under baseline cell culture conditions, we found overlapping metabolic features (alanine, aspartate, glutamine, arginine, proline, glycine, serine, and threonine) occurring across all three breast cancer cell lines studies, and metabolic signatures that were unique to specific cell lines (MCF7: sphingolipid and glycerophospholipid metabolism; MDA-MB-231: taurine and hypotaurine metabolism; HCC1937: glyoxylate and dicarboxylate metabolism) (**Figure 5-6**).

Our analysis of metabolites significantly altered in response to olaparib treatment correlate with reports from Bhute *et al*, where metabolic markers of PARP inhibition were reported as changes in amino acid metabolism (glutamine and alanine), downregulation of osmolyte levels (taurine, and GPC), phosphocreatine, lactate and pyruvate in MCF7 cells (28). We reported downregulation of those metabolites also in the MDA-MB-231 and HCC1937 cells, while low levels of fumarate were observed only in the HCC1937 cells (**Figure 6**). Bhute *et al.* also reported increased NAD+ levels for cells treated with veliparib. In our results NAD+ levels increased in the MCF7 cells treated with olaparib at the IC_10_ treatment concentration, accompanied by a decrease in NAD+ levels at ascending concentrations of olaparib. Reduced levels of NAD+ were also detected in the MDA-MB-231 and HCC1937 cells at all treatment concentrations. Recent studies have shown that in TNBC cells, olaparib enhances the signalling pathways of other NAD+-dependent deacetylase (i.e., sirtuins) (28, 29). These findings are in agreement with our observation of depleted acetyl-amino acid levels and enrichment of methyl-pyridines, -pyrrolidines, and -nucleosides. Further studies are needed to confirm the divergence of NAD+ flow towards alternative pathways and its association with specific breast cancer subphenotypes.

Glutamine- a precursor for protein, nucleotide, and lipid biosynthesis- is a fundamental amino acid in breast cancer cell metabolism, playing a pivotal role in providing anaplerotic intermediates for the tricarboxylic acid (TCA) cycle (30). Previous reports have indicated a reduction of glutamine levels only for the TNBC cells after treatment with veliparib, and in the MCF7 cells only in combination with other DDR inhibitors (16, 28). Our results show reduced glutamine levels in all cell lines treated with olaparib, suggesting increased glutamine utilisation. Once internalised by cells, glutamine can be converted to glutamate and alpha-ketoglutarate (α-KG). α-KG- a by-product of isocitrate- is oxidised in the TCA cycle through a reaction catalysed by isocitrate dehydrogenase (IDH), which is frequently mutated in cancer. Several studies have studied α-KG as an oncometabolite, where elevated levels induce the reversal of enhanced glycolysis through downregulation of the Hypoxia-inducible factor (HIF1), which following PARP inhibition leads to cell death (31, 32). Recent findings have shown that mutant IDH - and the consequent synthesis of aberrant α-KG forms - confers a BRCAness phenotype (33), downregulating the expression of the DNA repair enzyme Ataxia-telangiectasia mutated (ATM) kinase (34), altering the methylation status of loci surrounding DNA breaks (35). Together, these alterations lead to homology-dependent repair (HDR) impairment and increase susceptibility to PARP inhibition. On this basis, the reduced α-KG levels observed in olaparib-treated MDA-MB-231 and HCC1937 cells shows the basis for potential resistance to the anti-proliferative effects of olaparib. The increased utilisation of α-KG by HCC1937 cells, is paralleled by an increased consumption of serine at ascending doses of olaparib (**Table S5**). These observations are consistent with reports that in BRCA1- mutated TNBC cell lines, approximately 50% of α-KG results from the flux of serine metabolism (36).

Glutamine is also a source of nitrogen groups for the synthesis of nucleobases and nucleotides, either directly or through a process involving the transamination of glutamate and the TCA cycle-derived oxaloacetate that generates aspartate (37–39). Our results show low levels of glutamine are associated with overall reduction in nucleobase and nucleotide levels. MCF7 and HCC1937 cells showed accumulation of adenosine monophosphate (AMP), which represents a depleted energy and nutrient status of the cells known to activate the metabolic sensor AMP-activated protein kinase (AMPK) leading to cell growth inhibition (40). Different studies have considered activation of AMPK a metabolic cancer suppressor and an attractive therapeutic target for TNBC (41), however, its signalling network in response to PARP inhibition in different breast cancer cells needs to be established. Opposite to what observed by Bhute *et al*, Aspartate, a byproduct of the TCA cycle, accumulated in the MDA-MB-231 cells after PARP inhibition compared to its reduction in the MCF7 and HC1937 cells. Lowered plasma aspartate levels have been diagnosed in breast cancer patients suggesting an increased tumour utilisation of this metabolite (42). Moreover, we observed that aspartate metabolism is relevant both in the baseline model and in response to olaparib, which suggests a role of this metabolite in regulating the different metabolic phenotypes of breast cancer cells.

However, its role has been poorly investigated and little is known about its association with PARP inhibition. Among the pathways of aspartate utilisation, asparagine is converted through the enzyme asparagine synthetase (ASNS). The reaction requires glutamine as a substrate and consumption of adenosine triphosphate (ATP) to produce adenosine monophosphate (AMP) and pyrophosphate (PPi). Physiological levels of asparagine occur at levels of <0.05 mM in human plasma (44). Cancer cells harbouring mutant KRAS (e.g. MDA-MB-231), possess lower ASNS expression levels, leading to lower baseline aspartate levels explaining the rationale for the lack of aspartate detection in MDA-MB-231 lines (45). In breast cancer cells the increased bioavailability of asparagine promotes metastatic progression (45), due to its role in protein synthesis and regulation of amino acid homeostasis (46). We found elevated asparagine levels in olaparib-treated MCF7 and HCC1937 cells, suggesting a role for asparagine in the observed responses to exposure to PARP inhibitor.

Beyond asparagine synthesis, aspartate amidation through ASNS presents a source of amino building blocks for the synthesis of arginine in the urea cycle, which is in turn responsible for the synthesis of polyamines catalysed by ornithine decarboxylase (ODC). Polyamine accumulation previously has been correlated with the increased proliferation of both hormone-dependent and independent breast cancer cells (47), and recently found to contribute to BRCA1-mediated DNA repair (48). Moreover, metabolic profiling of plasma samples from patients with TNBC revealed an increase of diacetyl spermines associated with elevated expression of MYC, a well-known oncogene driving TNBC development and proliferation. Here, we found elevated diacetyl spermine levels following olaparib treatment in both TNBC and non-TNBC cells, suggesting an upregulation of polyamine catabolism, irrespective of cell line BRCA- and hormone receptor-status. Parallel to their relevance in cellular metabolism, amino acids serve also as biological buffers through regulation of cellular pH. Low extracellular pH is associated with positively charged amino acids and a known hallmark of cancer arising from enhanced glycolysis, production and altered lactate metabolism, resulting in altered mTOR pathway activation, ultimately regulating cancer cell metabolism (49, 50).

Glutathione (GSH), is involved in the protection against ROS and regulation of intracellular redox homeostasis. Elevated GSH levels have previously been reported in TNBC compared to luminal breast cancers, suggesting the relevance of GSH to our observations of lower sensitivity to olaparib in TNBC cell lines (17, 51).

Lipids mediate various cellular biological functions, including energy storage, cell membrane structural composition and signal transduction, the increased biosynthesis of which is a marker of metabolic rewiring observed in malignant breast cancers (52, 53). Our findings show downregulation of fatty acid biosynthesis following olaparib treatment, with a reduction in phospholipid levels including lysophosphatidylcholines and glycerolphosphocholines in all cell lines. Poly-unsaturated fatty acids (PUFAs), have previously been implicated in MCF7 and MDA-MB-231 cell apoptosis through the induction of lipid peroxidation and altered cellular redox state (54). Moreover, elevated PUFA levels have been associated with the proteolytic cleavage of PARP and its inhibition, leading to cell death (55). On this basis, the reduced PUFA levels observed in HCC1937 cells may indicate their resistance to olaparib treatment. Only a limited number of studies have reported a correlation between PUFAs and breast cancer subphenotypes, requiring further validation by additional studies.

Future targeted metabolomics studies using additional TNBC cell lines and clinical tumour clinical specimens are required to validate our observations. Validation of our findings could define prognostic biomarkers that will aid diagnose and enable the implementation of precision medicine in the management of breast cancer.

## CONCLUSION

Our data show differential sensitivity of breast cancer cell lines to olaparib treatment that was dose-dependent and demonstrated the increased sensitivity of TNBC cells to DNA damage foci accumulation. The application of metabolomics to the study of breast cancer remains in its infancy, with only a handful of studies reporting combined metabolomics and phenotypic analyses. Data acquired from metabolomics analysis can be validated against routine molecular biology and phenotypic assays, providing a powerful platform for biomarker detection or the discovery of novel actionable pathways for drug development.

Our results show that fingerprinting the metabolic profile of cells can be a powerful tool for uncovering potential oncometabolites or mechanisms giving rise to chemoresistance. Findings from such studies may provide potential additional actionable targets for modulating response to drug treatment or the design of new drug combinations that will overall enhance DNA damage efficacy, ultimately improving patient response to radiotherapy and adjuvant chemotherapy.

## ACKNOWLEDGEMENTS

The authors acknowledge funding of these studies from Tenovus Scotland and the Royal Society of Edinburgh Research Reboot funding. We acknowledge support from the Strathclyde Centre for Molecular Biology (SCMB) for providing access to mass spectrometry facilities. ZR acknowledges funding from the UK Engineering and Physical Sciences Research Council (EPSRC-EP/V028960/1).

## AUTHOR CONTRIBUTIONS

Conceptualization: ZR, NJWR, and DB.; Investigation: DB, YH, GF, NJWR, ZR.; Methodology: DB, YH, LvDD, GF, ZR, NJWR.; Analysis: DB, YH, NJWR, ZR, LvDD.; Writing original draft: ZR and DB.; Visualization: DB, LvDD, NJWR, ZR.; Writing- reviewing and editing: DB, YH, LvDD, GF, NJWR, ZR.; Funding acquisition: NJWR and ZR. All authors have read and agreed to the published version of the manuscript.

## DATA AVAILABILITY STATEMENT

The datasets generated and used/or analysed are available from the corresponding authors upon request.

## CONFLICTS OF INTEREST

The authors declare no conflict of interest.

## Supplementary Information

**Table S 1.** Cell lines used in this study and their corresponding clinicopathologic profiles (ER: estrogen receptor, PR: progesterone receptor, and HER2: Human epidermal growth factor 2 receptor)

| Cell Line | Histology | Subtype | Immunoprofile | Genetic alterations |
| --- | --- | --- | --- | --- |
| **MCF-7** | Metastatic Adenocarcinoma | Luminal A | ER+, PR+, HER2- | PIK3CA, CDKN2A, GATA3, PIK3CA, TP53 |
| **MDA-MB-231** | Metastatic Adenocarcinoma | Basal | ER-, PR-, HER2- | BRAF, CDKN2A, KRAS, NF2, TP53 |
| **HCC1937** | Primary Ductalcarcinoma | Basal-like | ER-, PR-, HER2- | BRCA1, TP53 |

### Elution Gradient used for LC-MS

**Buffer A composition:**10 mM ammonium acetate in 95% acetonitrile, 5% water with 0.1% acetic acid

**Buffer B composition:** 10 mM ammonium acetate in 50% acetonitrile, 50% water with 0.1% acetic acid

**Table S2.**
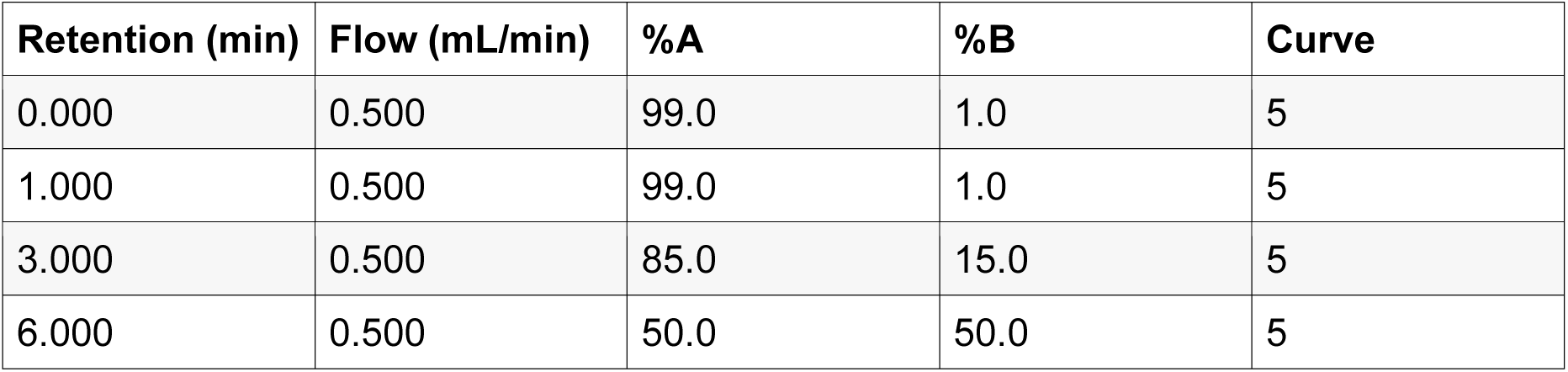
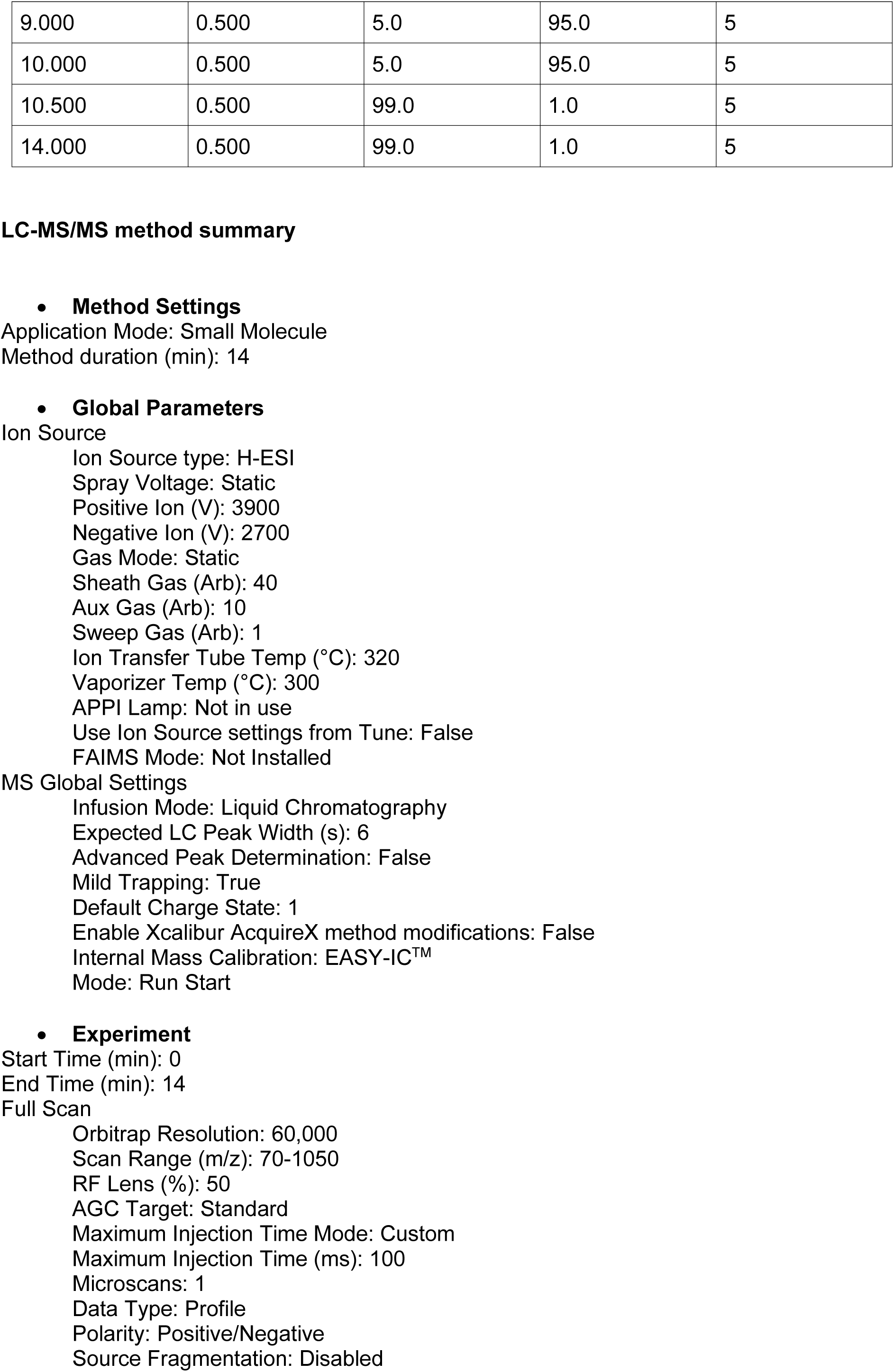
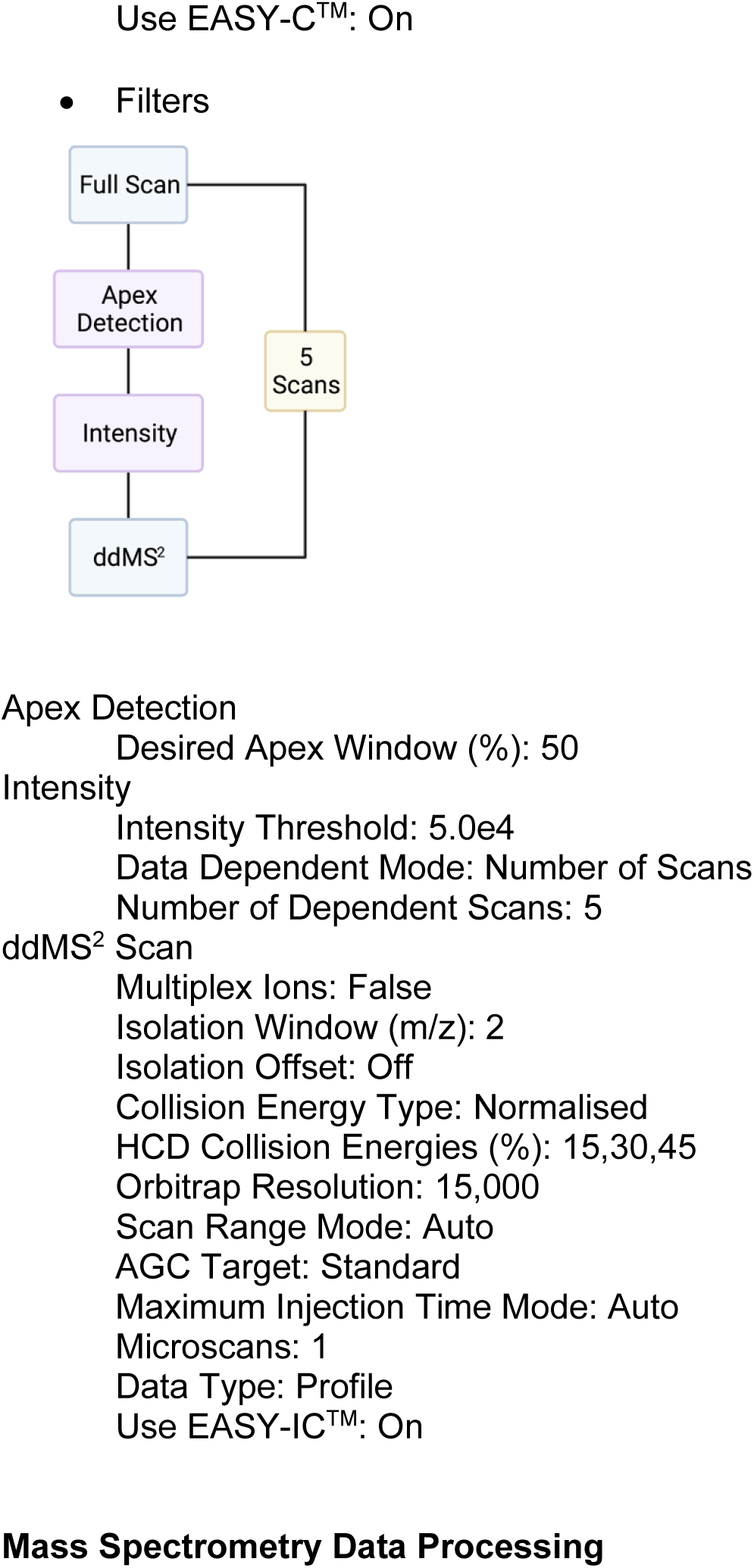
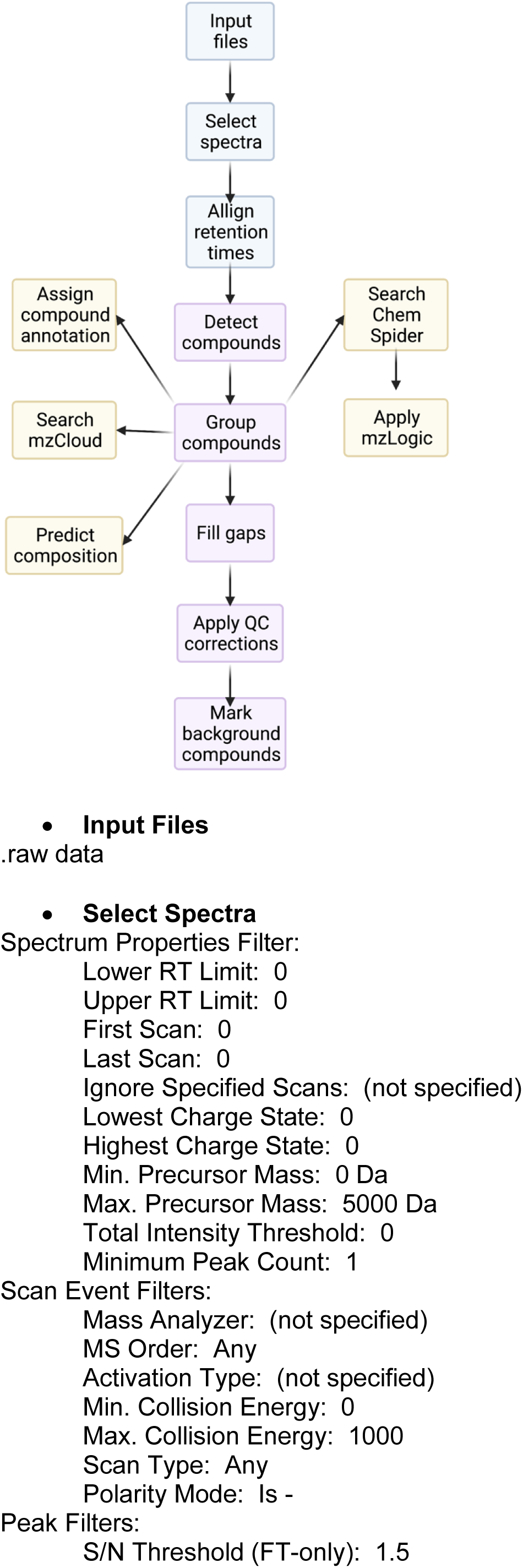
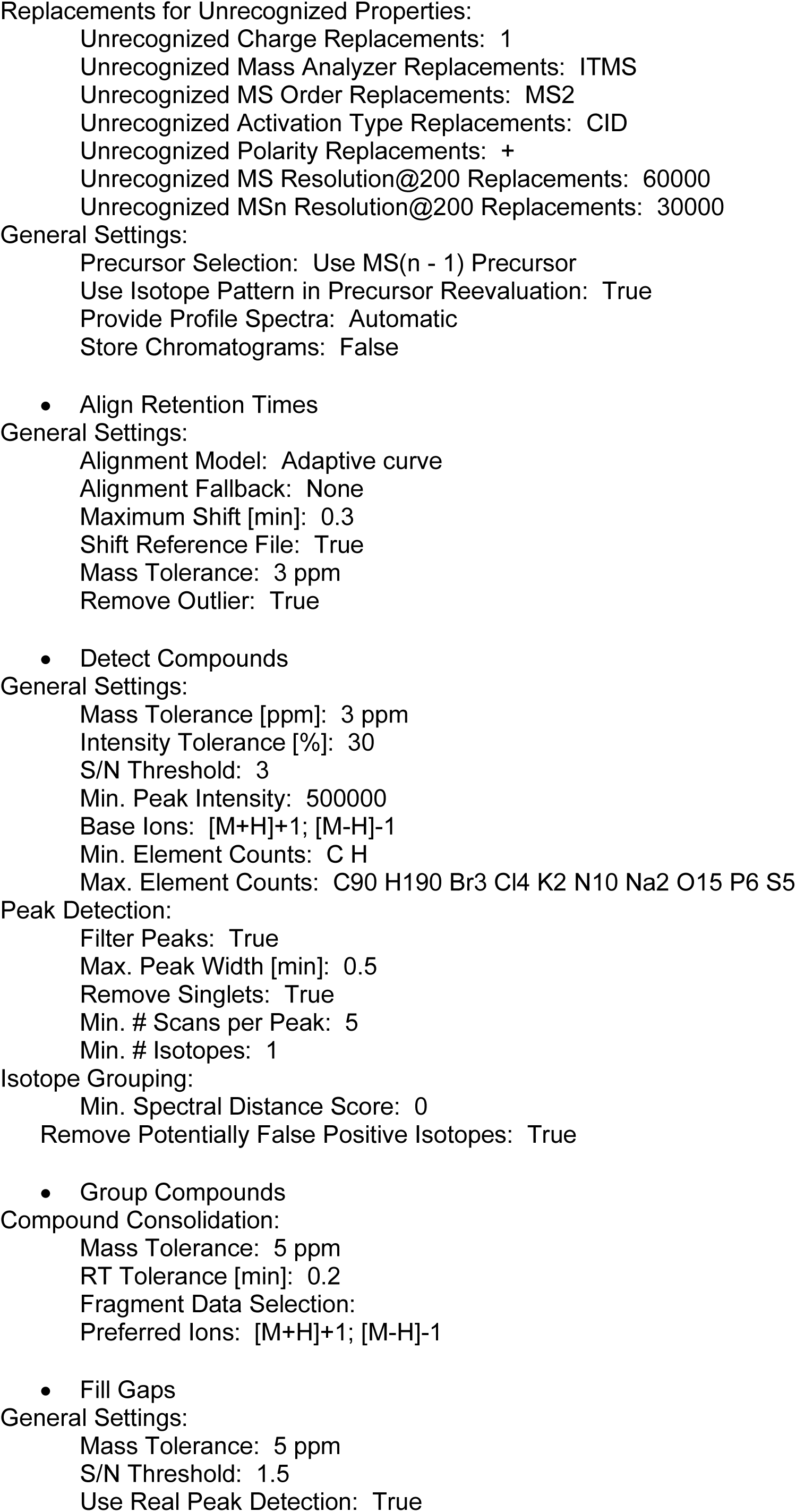
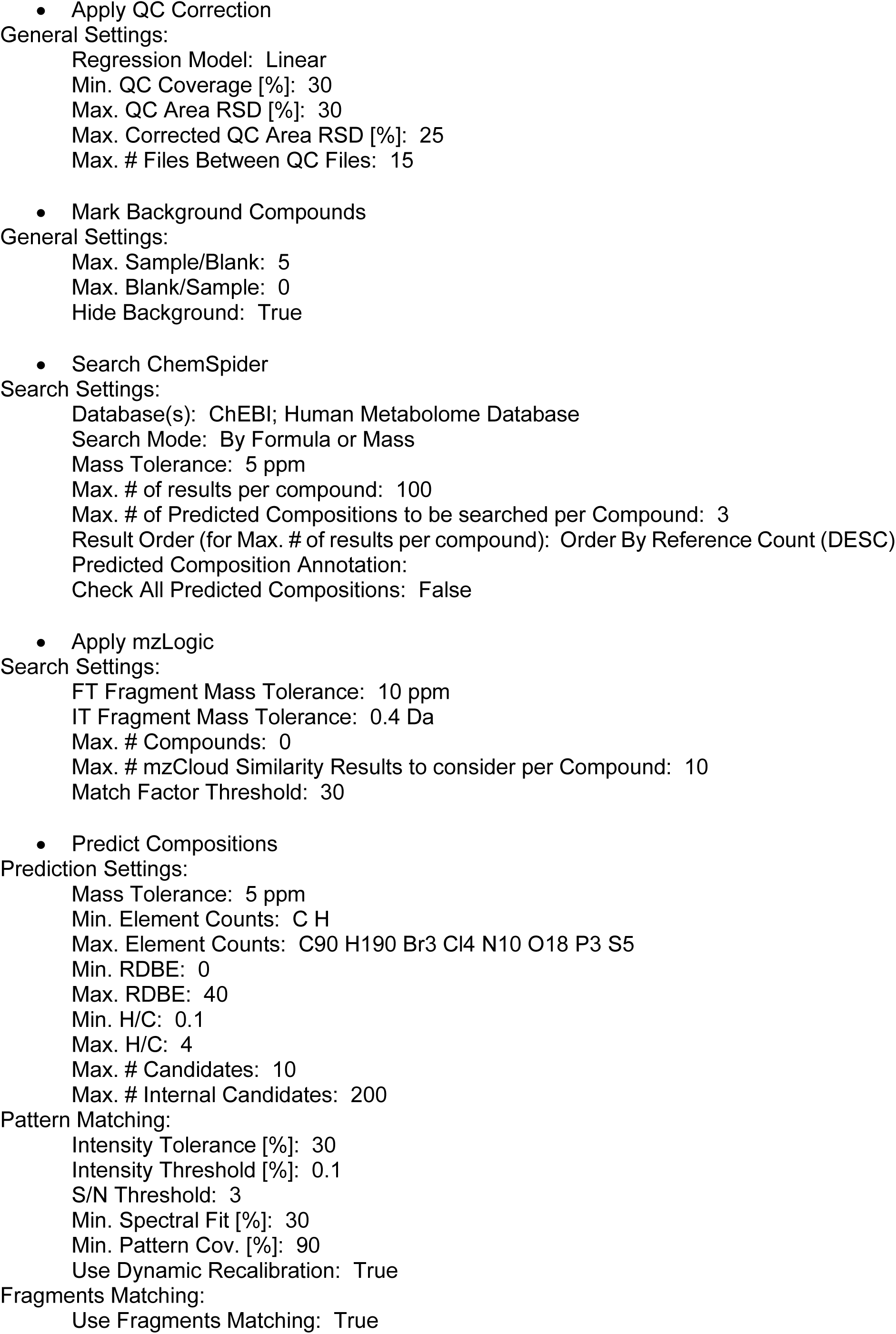
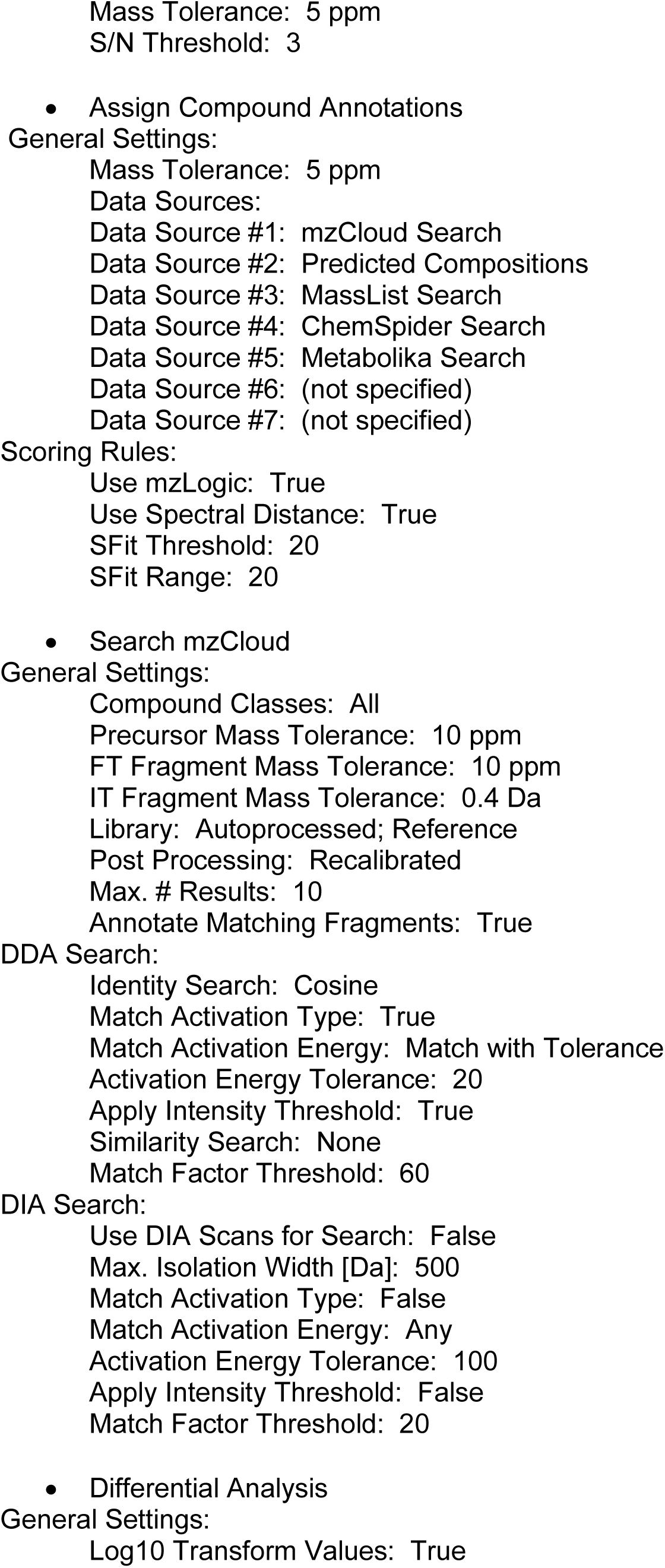
Corresponding elution gradient used for the chromatographic separation of metabolite extracts

**Table S3.**
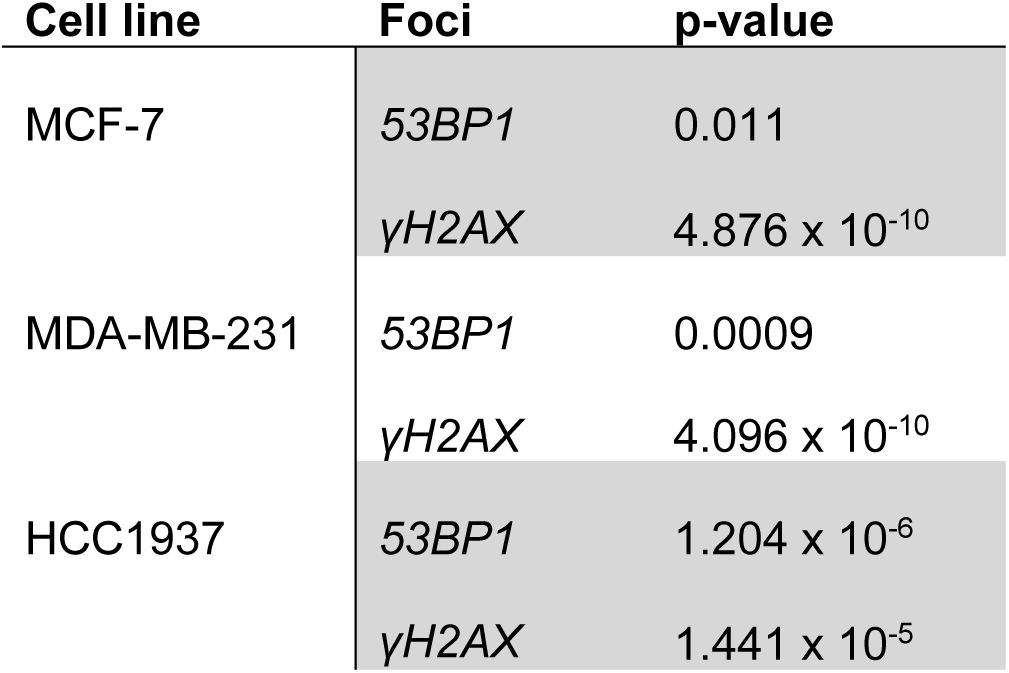
ANOVA analysis of olaparib dose-dependent DNA DSB immunofoci formation

| Cell line | Foci | p-value |
| --- | --- | --- |
| MCF-7 | 53BP1 | 0.011 |
| | γH2AX | $4.876 \times 10^{-10}$ |
| MDA-MB-231 | 53BP1 | 0.0009 |
| | γH2AX | $4.096 \times 10^{-10}$ |
| HCC1937 | 53BP1 | $1.204 \times 10^{-6}$ |
| | γH2AX | $1.441 \times 10^{-5}$ |

**Panel of individual PCA pairwise analysis of MCF7, MDA-MB-231 and HCC1937 at their relative IC_10_, IC_25_ and IC_50_ doses of Olaparib**

**Figure S 1.**
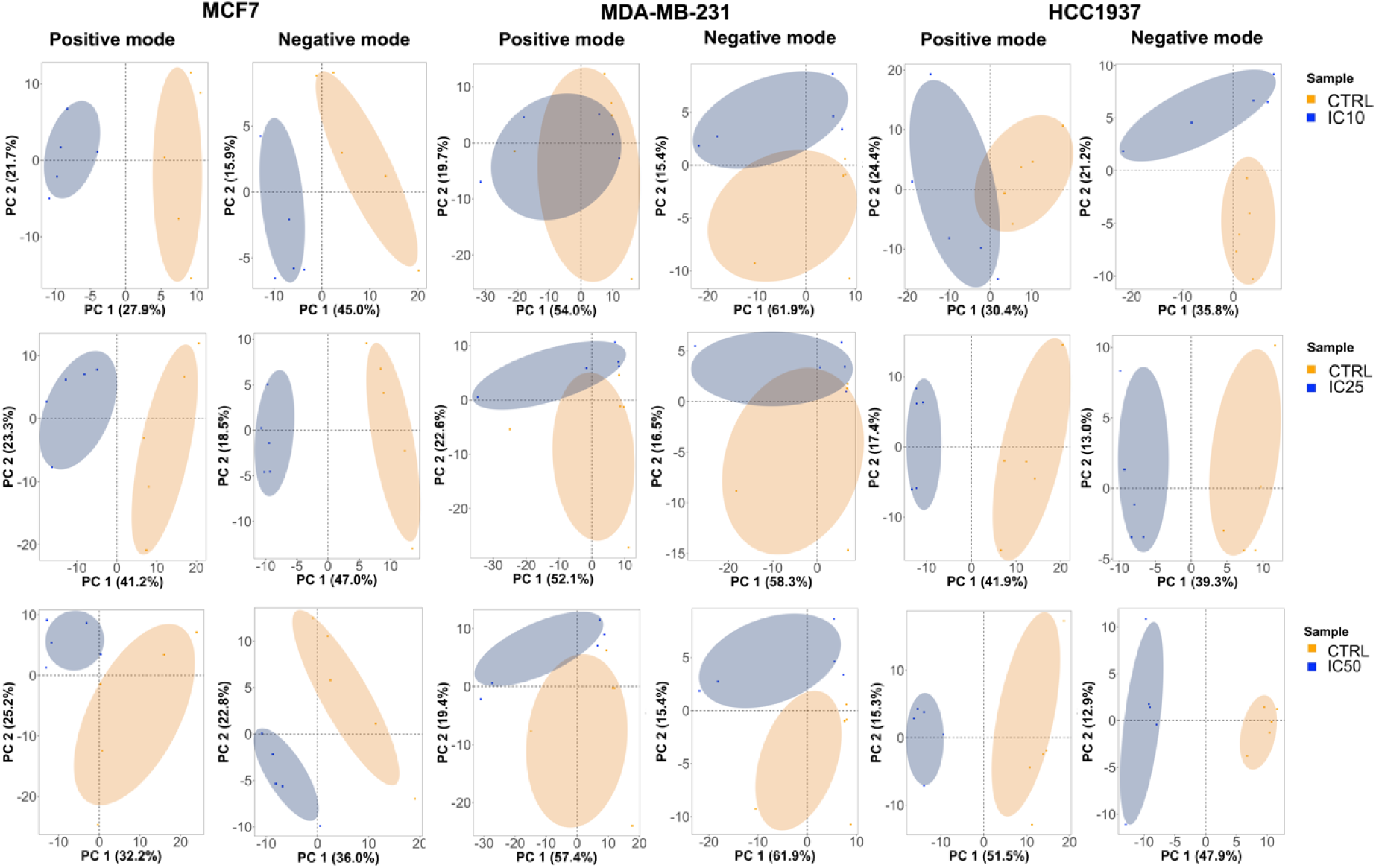
PCA pairwise analysis of untargeted metabolomics data collected, in both positive and negative mode, from MCF7, MDA-MB-231, and HCC1937 cells treated with IC_10_, IC_25_ and IC_50_ olaparib treatment doses. Data points in the two-dimensional PCA score plot were central scaled. The plot was designed on R through the ggplot2 graphical package (n=5).

**Pairwise differential analysis of metabolites identified in MCF7, MDA-MB-231 and HCC1937 cells in positive and negative mode**

**Figure S2.**
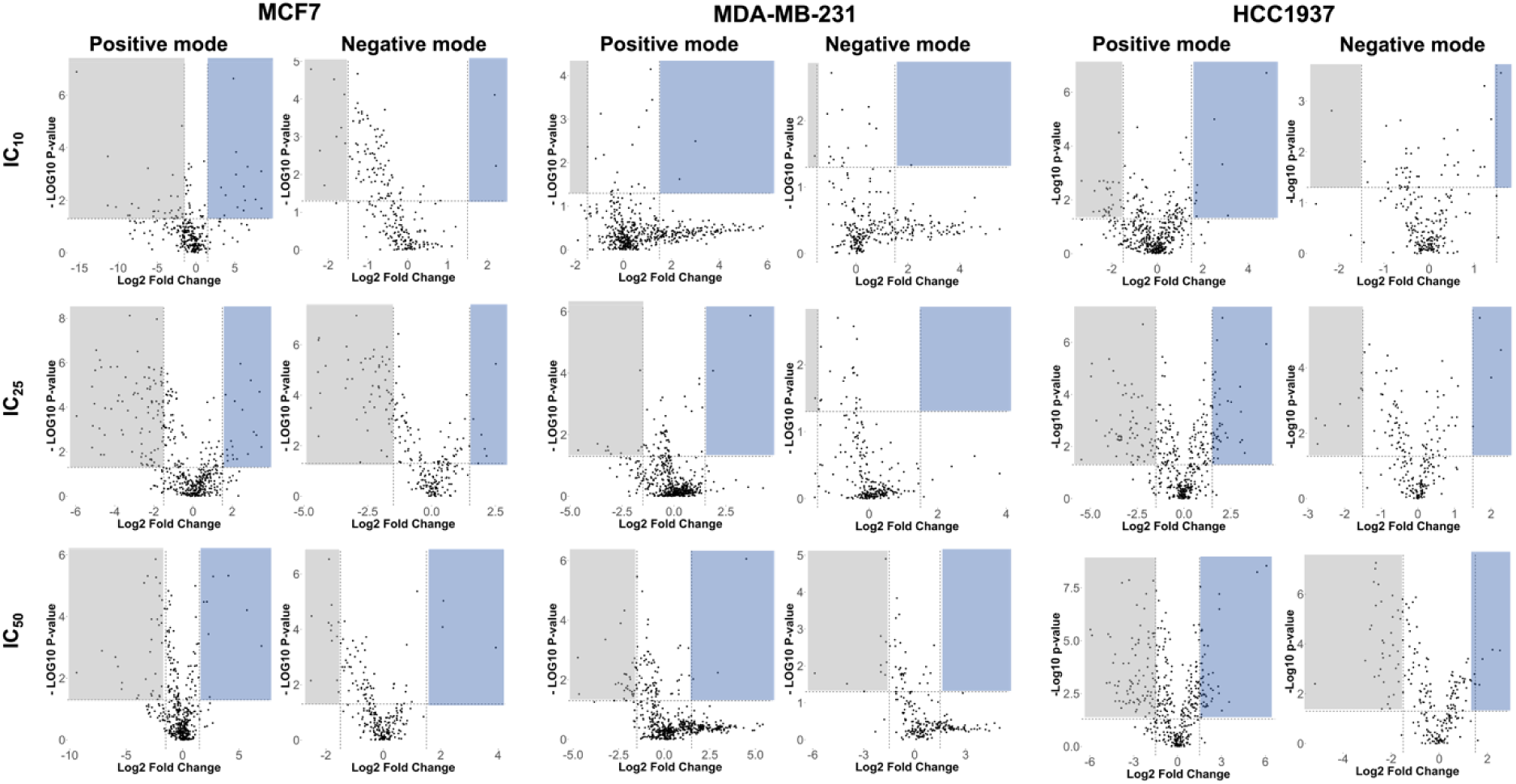
Volcano plots showing the log2 fold change and the -log10 adjusted p-values in metabolite levels induced by treatment with different doses of Olaparib (IC_10_, IC_25_, and IC_50_) in MCF7, MDA-MB-231 and HCC1937 cells. Data were selected at the cut off values adjusted p<0.05 and fold change >1.5. Blue and grey boxes indicate metabolites significantly enriched and depleted in the different samples, respectively.

**Table S4.** Global differential number of altered metabolites for samples treated with IC_10_, IC25 and IC_50_ of Olaparib and their relative control (non-treated) samples. Metabolites identified in both positive and negative mode with p-value = <0.05 and Log2 Fold Change = >1.5.

| Sample | HESI + | HESI - |
| --- | --- | --- |
| MCF7 IC10/Ctrl | 41 | 10 |
| MCF7 IC25/Ctrl | 111 | 62 |
| MCF7 IC50/Ctrl | 41 | 15 |
| MDA231 IC10/Ctrl | 2 | 1 |
| MDA231 IC25/Ctrl | 12 | 1 |
| MDA231 IC50/Ctrl | 34 | 9 |
| HCC1937 IC10/Ctrl | 36 | 2 |
| HCC1937 IC25/Ctrl | 107 | 13 |
| HCC1937 IC50/Ctrl | 134 | 43 |

**Figure S3.**
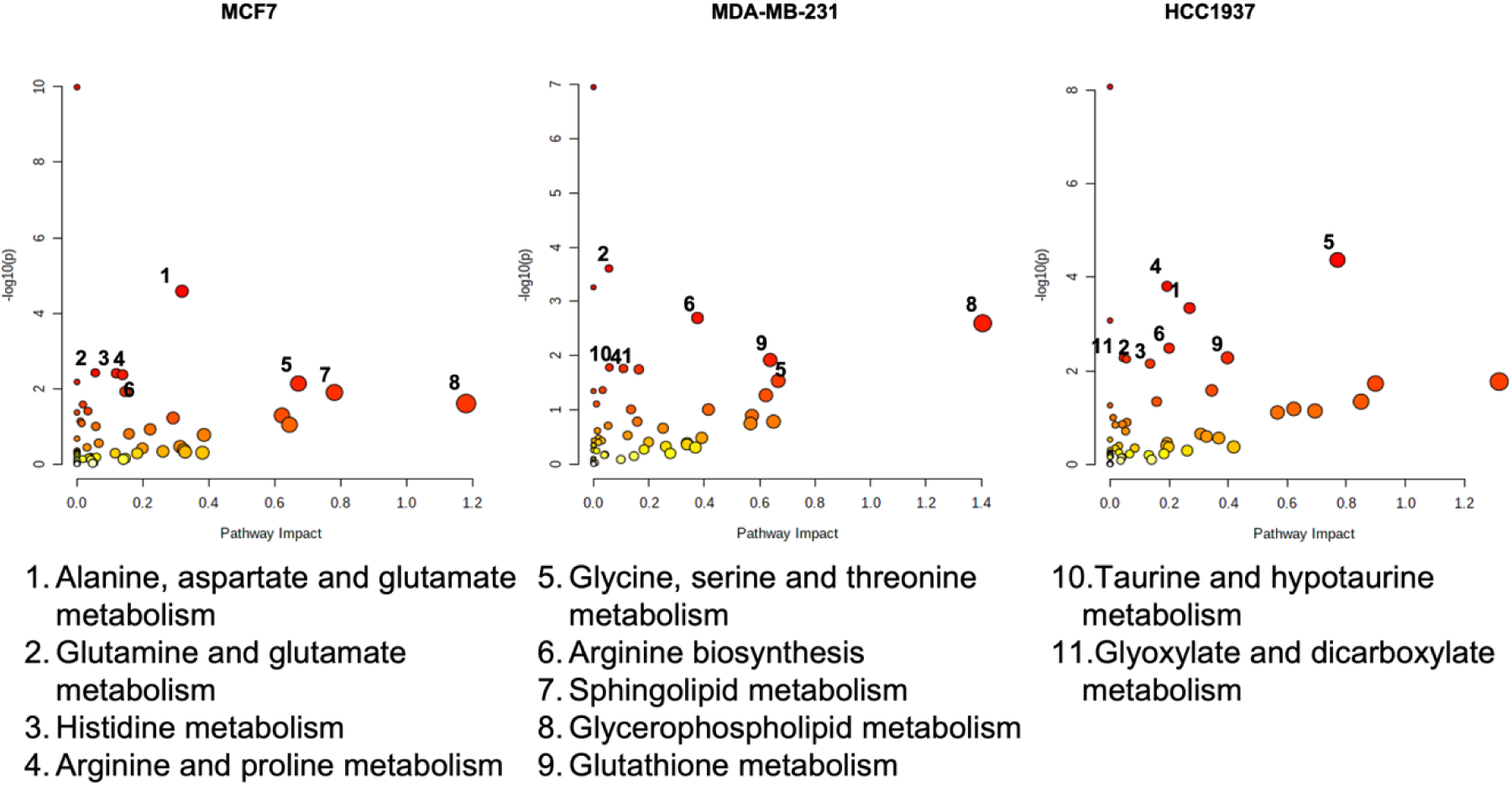
Enrichment analysis of non-treated MCF7, MDA-MB-231 and HCC1937 cells.

**Table S5.**
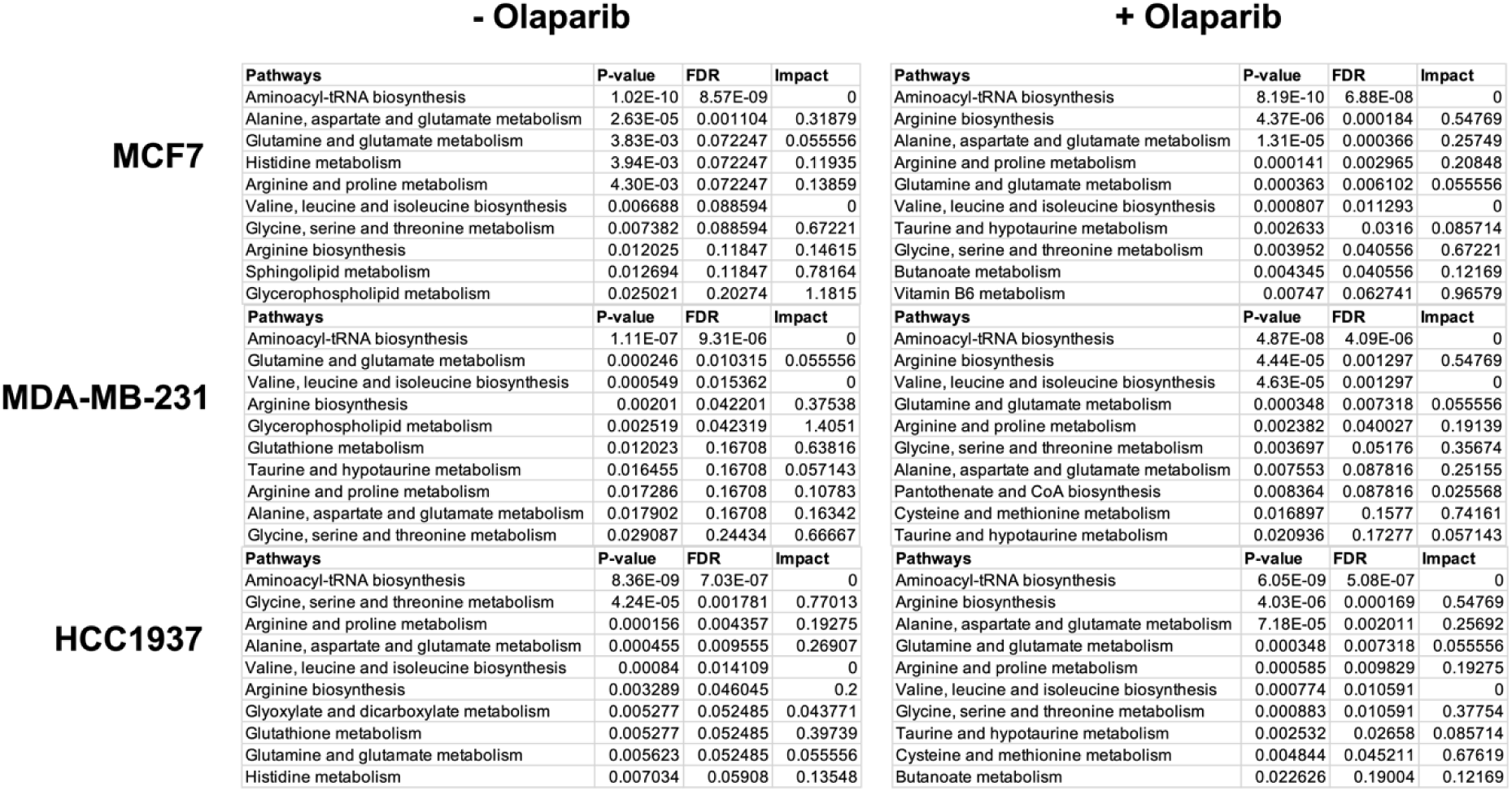
Metabolic pathways in different breast cancer cells (MCF7, MDA-MB-231, and HCC1937) before and after treatment with IC_50_ dose of Olaparib. FDR = False Discovery Rate.

**Figure S4.**
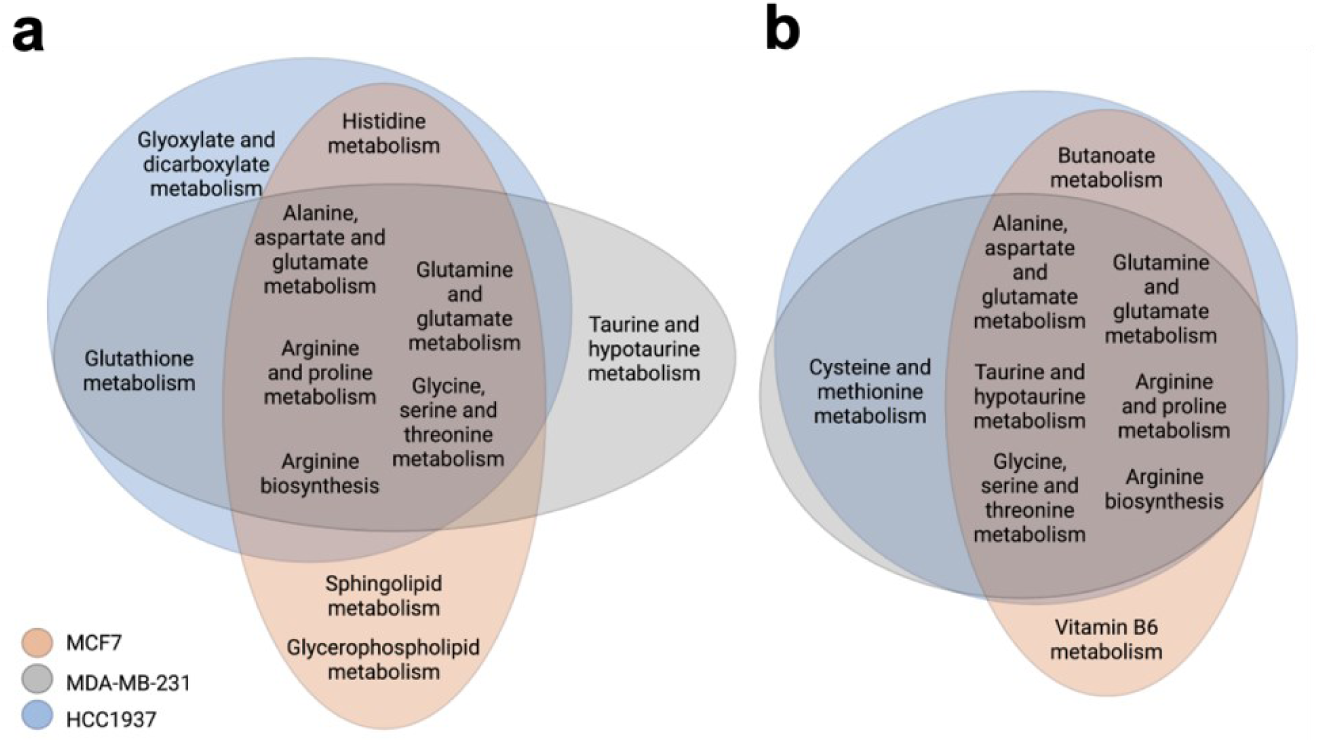
Venn diagram representing the metabolic pathways in MCF7, MDA-MB-231 and HCC1937 cells .a) Baseline metabolic pathways and b) following a seven day treatment with olaparib at IC_50_ doses.

**Table S6.**
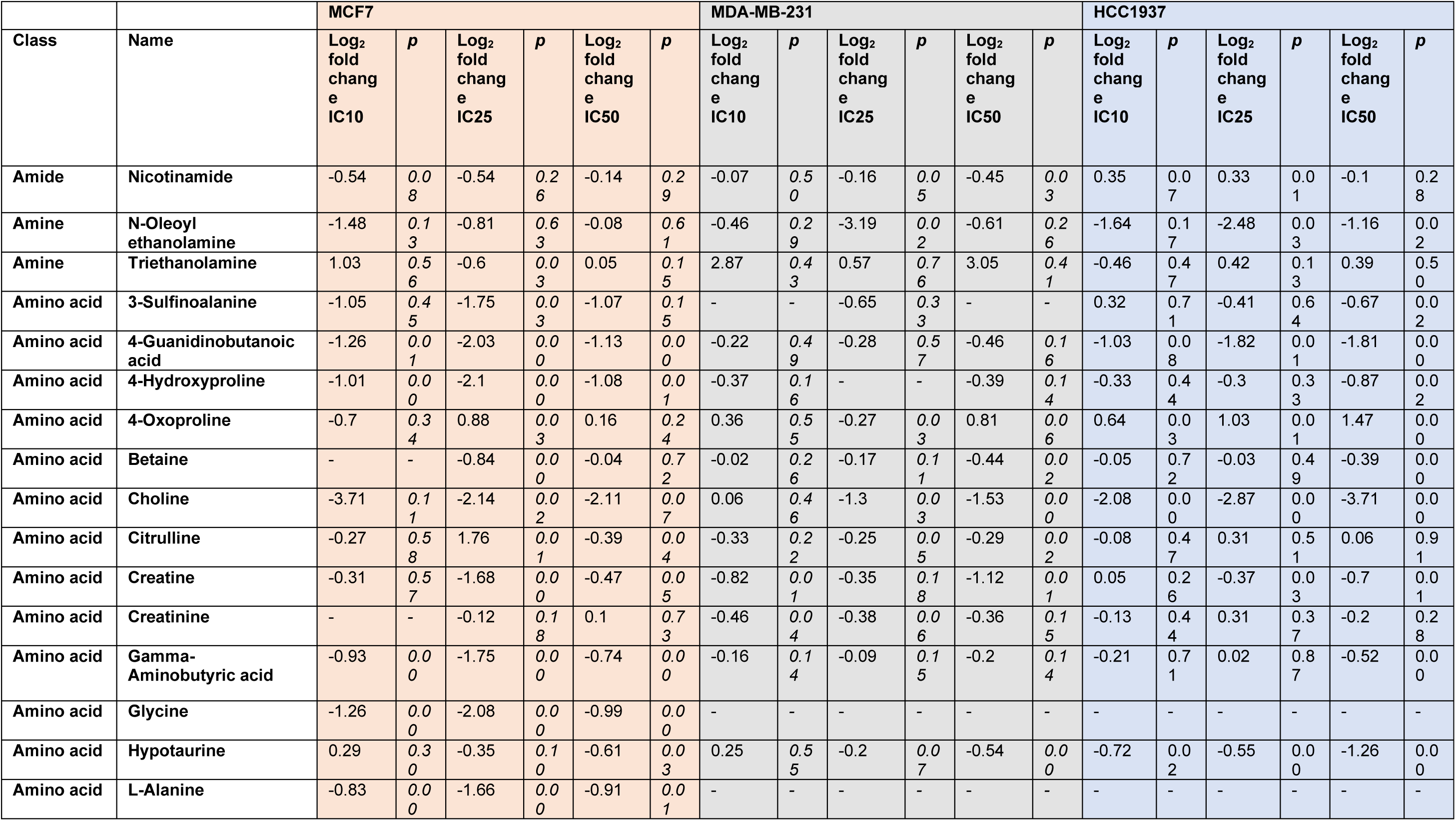
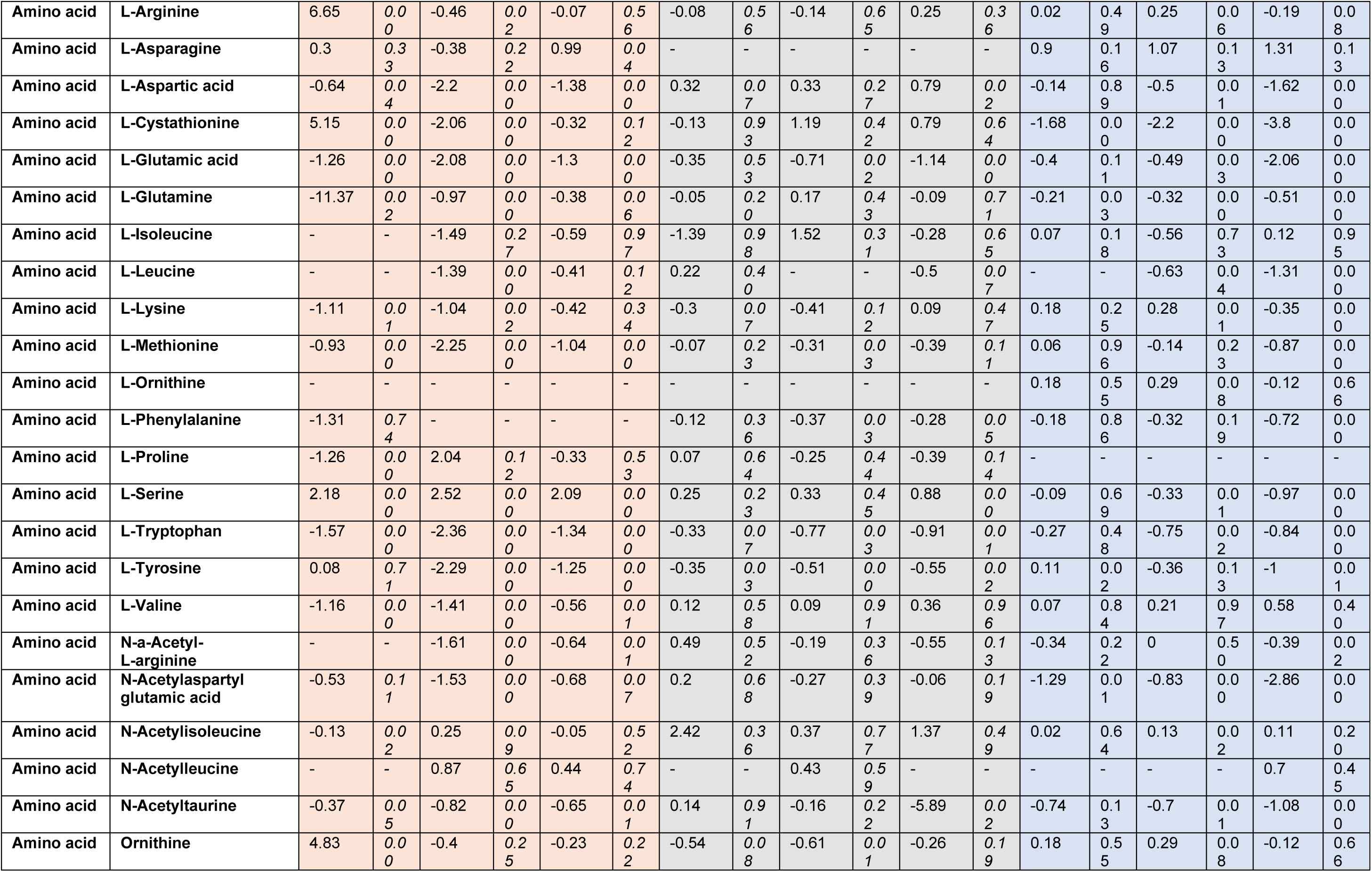
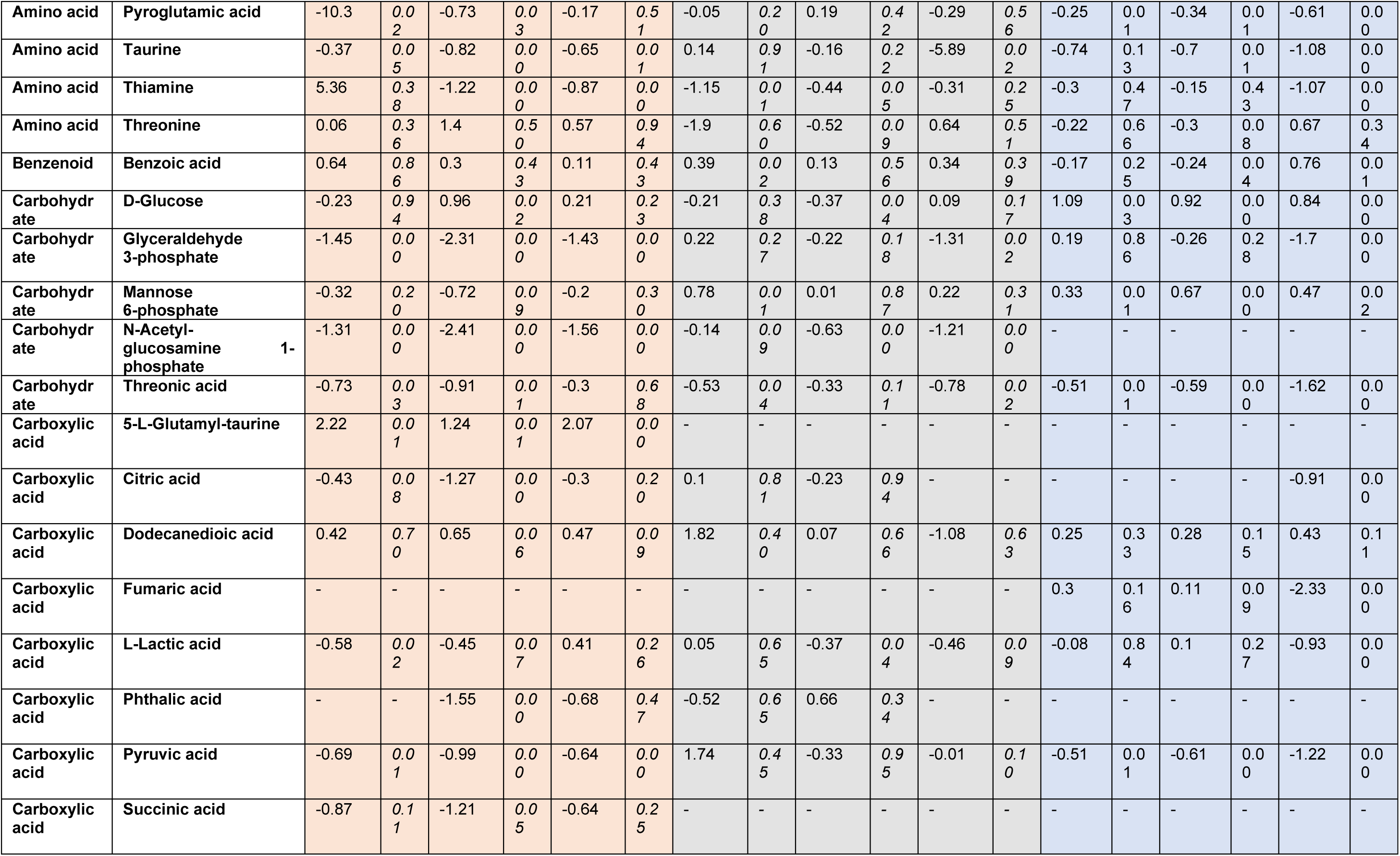
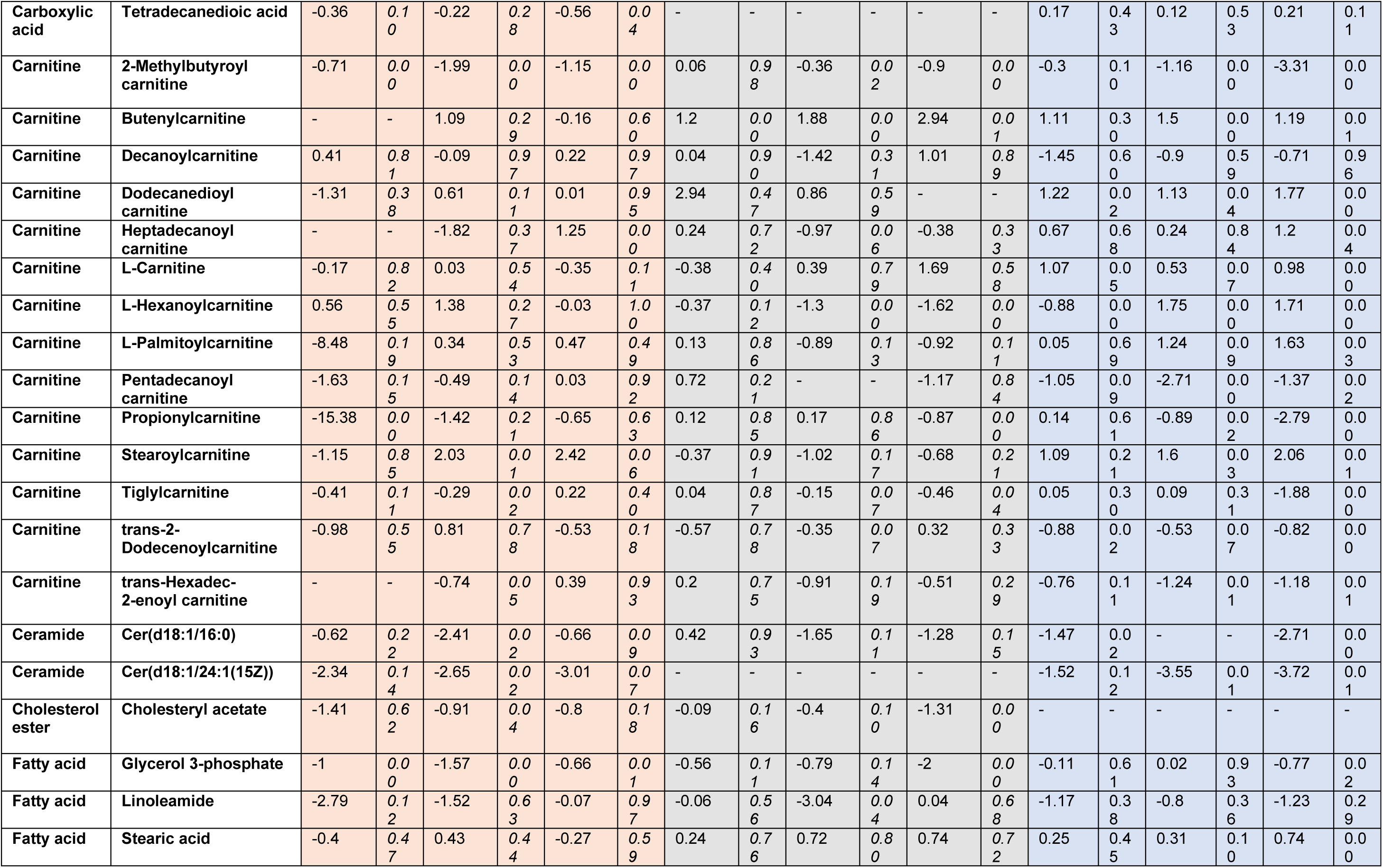
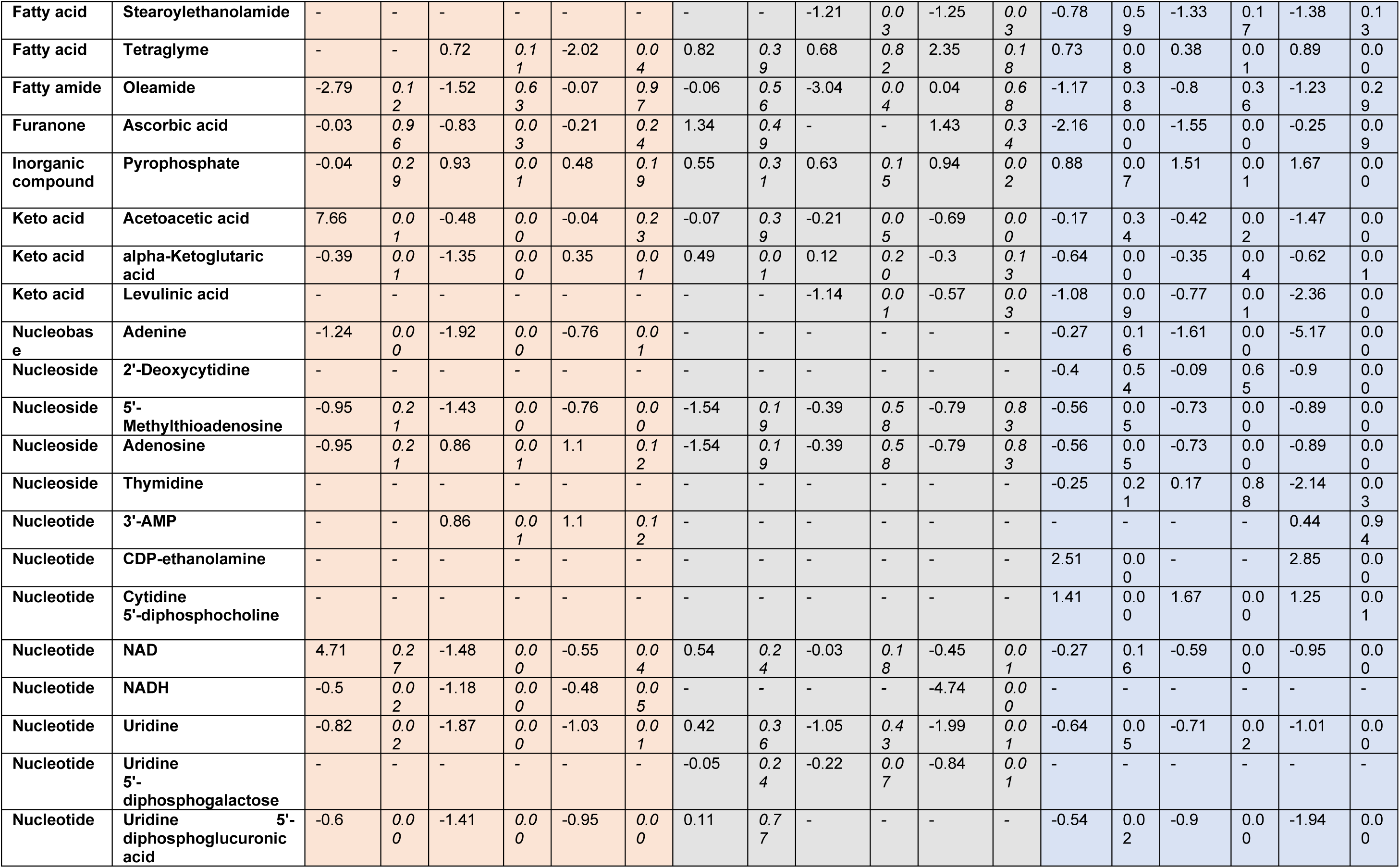
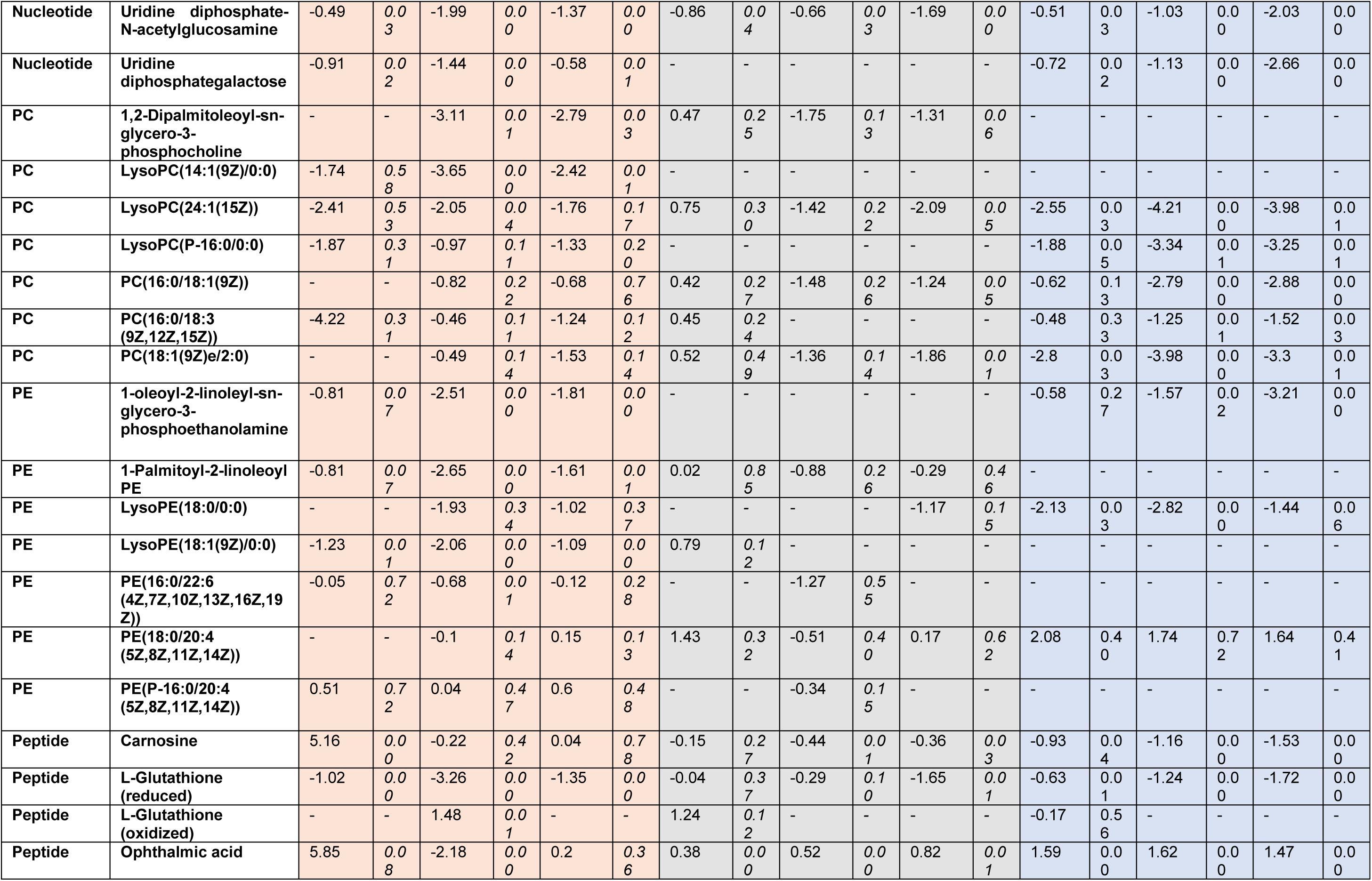
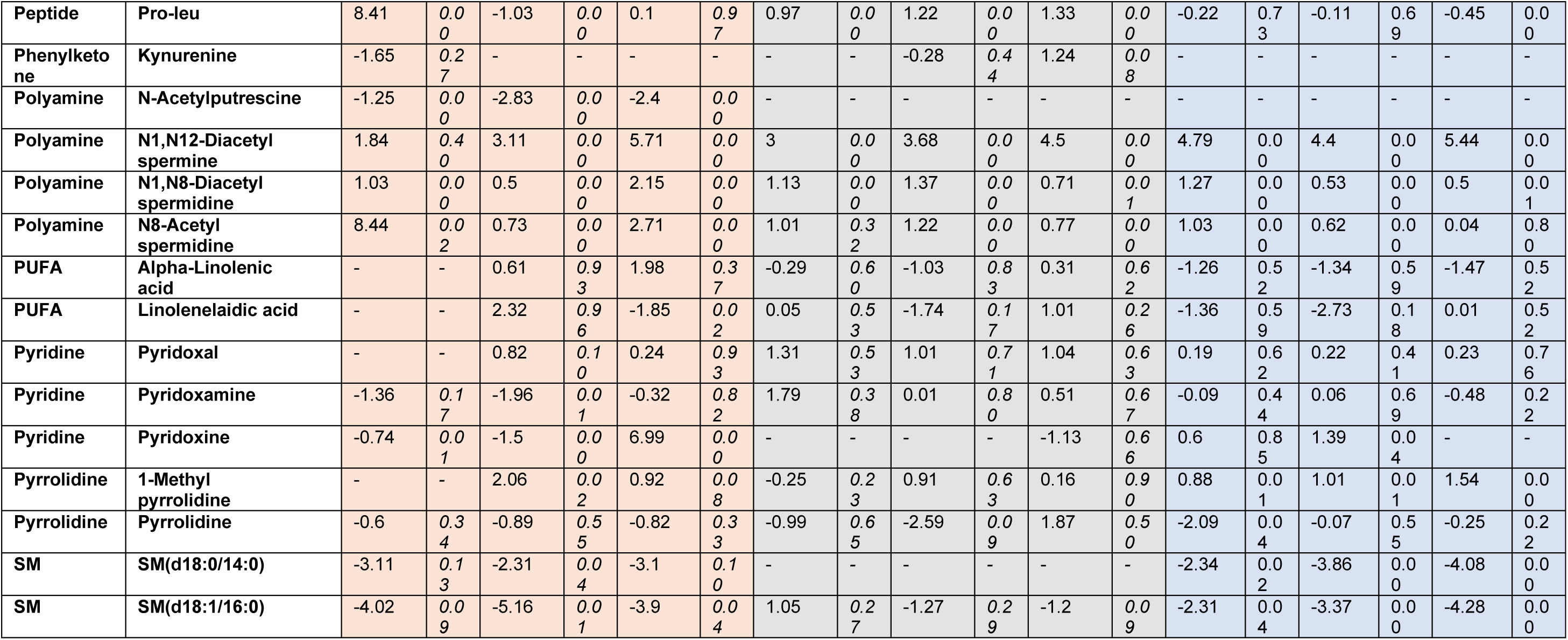
Classification of the metabolites identified in MCF7, MDA-MB-231 and HCC1937 at all Olaparib doses (IC_10_, IC_25_ and IC_50_) after seven days treatment. Class, name, Log_2_ fold change, and the p-value (p) is represented for each compound. PC: phosphocholine; PE: phosphoethanolamine; PUFA: poly unsaturated fatty acid; SM: Sfingomyelin.

**Figure S5.**
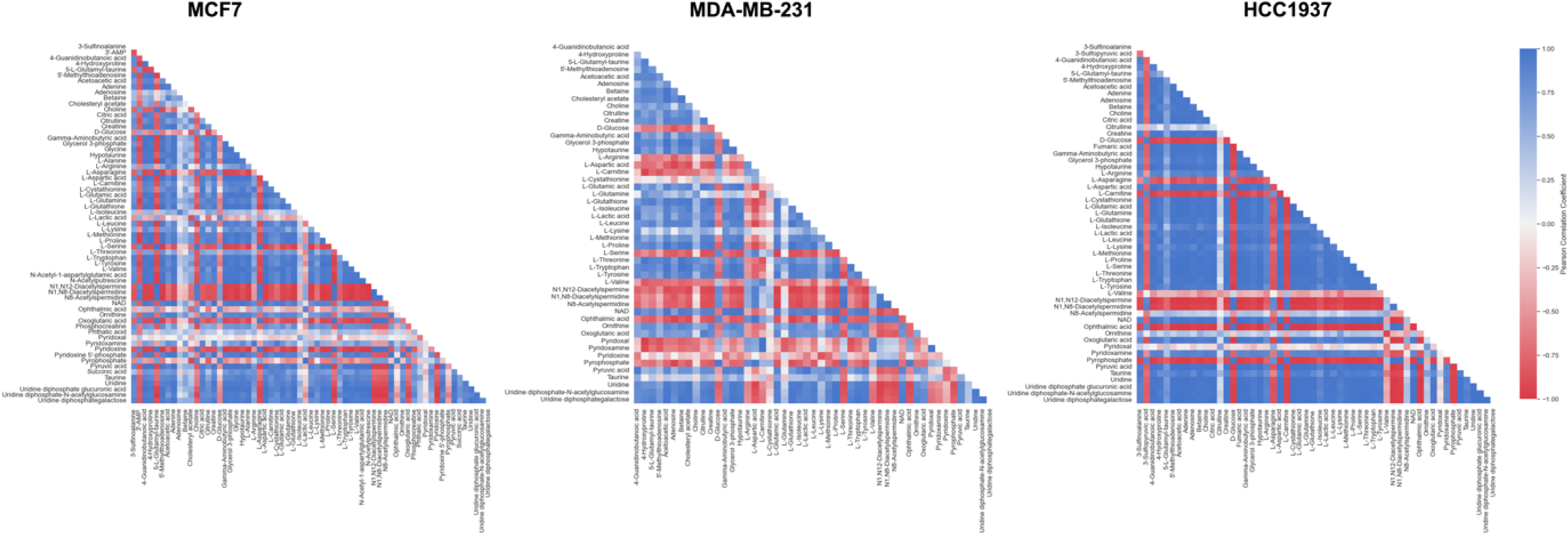
Pearson’s correlation analysis between the relevant metabolites identified within each different breast cancer cell line. Pearson’s coefficient is set in a range of 1 to -1, indicative of a positive and negative correlation, respectively.

## Notes

### Competing Interest Statement

The authors have declared no competing interest.

### Summary of Updates

Corrected figure numbers

